# DEC1 is an adult clock regulator that links circadian rhythms to functional β-cell maturation

**DOI:** 10.1101/2023.08.11.552890

**Authors:** Zihan Gao, Mai Liu, Ana P. Montalvo Landivar, Zoe L. Gruskin, Andrew Leduc, Yu Xie, Sam Preza, Jingyi Du, Xiaolei Liu, Victoria Huhn, Zhihui Cheng, Christopher Krapp, Matthew Haemmerle, Andrea V. Rozo, Xinyu Wen, Stephen J Lee, Stacy Liang, Akshaya Biju, June H. Ahn, Li Li, Juerg R. Straubhaar, Monica M. Cho, Wei Qin, Luisa Lopez Monzon, Chengyang Liu, Ali Naji, Rajan Jain, Nicolai Doliba, Doris A. Stoffers, Romina Bevacqua, Peter Klein, Nikolai Slavov, Juan R. Alvarez-Dominguez

**Author notes:** These authors contributed equally to this work (Zihan Gao, Mai Liu, Ana P. Montalvo Landivar).

## Abstract

Circadian clocks attune metabolism to daily energy cycles, but how they regulate functional maturation of metabolic tissues is poorly understood. Here we show that DEC1, a clock transcription factor induced in adult islet β cells, coordinates their glucose responsiveness by synchronizing energetic and secretory rhythms. DEC1 binds and regulates maturity-linked genes to integrate insulin exocytosis with β-cell energy metabolism, and β-cell-specific *Dec1* ablation disrupts their transcription synchrony. *Dec1*-disrupted mice develop lifelong glucose intolerance and insulin deficiency, despite normal islet formation. Metabolic dysfunction upon β-cell *Dec1* loss stems from poor coupling of islet insulin secretion to glucose metabolism, reminiscent of fetal/neonatal islet immaturity. We link stunted maturation to a deficit in circadian bioenergetics, prompted by compromised glucose utilization, mitochondrial dynamics, and respiratory metabolism, which is rescued by increased metabolic flux. Thus, DEC1 links circadian clockwork to β-cell metabolic maturation, revealing a hierarchy for how the clock programs metabolic tissue specialization.

## Main Text

Optimizing energy use to fuel availability is critical for metabolic fitness^1,2^. Circadian clocks evolved to anticipate 24-hour cycles in energy availability tied to earth’s rotation^3,4^. In mammals, clocks throughout the body optimize critical energy-requiring tasks—from gene expression to protein synthesis—to daily energy cycles by programming cell-autonomous 24-hour rhythms^5,6^. These circadian rhythms emerge as cells differentiate along specific lineages^7,8^, yet how they regulate differentiated cells becoming functionally specialized, or “mature”, is not understood^9,10^.

The circadian clock mechanism comprises a core feedback loop between transcriptional activators (CLOCK/BMAL1) and repressors (PERs/CRYs), along with interlocking regulatory loops involving the ROR, REV-ERB, and DEC transcription factors^11^. These clock regulators play general and specific roles in metabolism, and metabolic effectors in turn provide regulatory feedback to the clock^5,12–14^. As a result, circadian clockwork shapes and is shaped by myriad metabolic functions—from nutrient uptake and transport to breakdown and storage. Accordingly, clockwork disruption or circadian misalignment fuel obesity, metabolic syndrome, and diabetes in animal models^15–17^ and humans^5,18,19^. However, major questions remain as to how the core and interlocking clock regulators coordinate metabolic shifts to enable a mature physiology^6,20^.

Pancreatic islet β cells, vital controllers of glucose homeostasis, offer a model to dissect how the molecular clock steers metabolic specialization. Mature β-cell function develops after birth, as glucose-coupled insulin secretion emerges and both the glucose threshold for secretion and secretion capacity increase^21–23^. Postnatal maturation co-occurs with emergence of the β cell-autonomous clock, which optimizes glucose-stimulated insulin secretion (GSIS) responsiveness to the active, feeding phase of the day^24–26^. Critically, *Clock*-mutant or *Bmal1*-ablated mice show stunted islet growth and GSIS responsiveness, leading to hypo insulinemic diabetes^17,27,28^.

Although the underlying mechanisms are unclear, the CLOCK/BMAL1 complex has been proposed to modulate GSIS responsiveness by binding to and programming rhythmic transcription of late secretory machinery^29^ and of the redox homeostasis regulator *Nrf2*^30^.

To elucidate the coordination of β-cell maturation by the circadian network, we focused on the transcription factor DEC1 (so-called as it is found in differentiated embryo chondrocytes; also called BHLHE40, SHARP2, STRA13, and BHLHB2). DEC1 is known to entrain circadian rhythms to environmental cues, including light and feeding, by competing with CLOCK:BMAL1 for DNA E-box binding and/or binding to BMAL1^13,14,31,32^. Roles for DEC1 have been described in regulating circadian entrainment, sleep, memory consolidation, and immune cell function^32–35^. However, DEC1’s role in β-cell development remains unexplored. We previously predicted that DEC1 partakes in the core regulatory circuit that defines mature β-cell identity^36^, leading us to hypothesize that DEC1 regulates β-cell metabolic maturation.

Here, we define DEC1’s targets in β cells and use genetic, single-cell, and biochemical studies to reveal its role in the daily control of insulin secretion and metabolic homeostasis. We find that DEC1 (but not CLOCK/BMAL1) binds and synchronizes the rhythmic transcription of maturity-linked genes integrating energy metabolism and GSIS. *Dec1*-ablated mice are hypo-metabolic and develop lifelong glucose intolerance and insulin deficiency, which we link to immature islet function resulting from poor coupling of insulin secretion to glucose stimulation. Single-cell DNA accessibility, RNA expression, and proteome profiling of *Dec1*-ablated β cells trace this defect to disrupted expression of regulators or effectors of insulin responses to glucose sensing and metabolism. Accordingly, circadian rhythms in glucose import, utilization, and mitochondrial flux are impaired in *Dec1*-ablated islets, whereas pharmacologic or genetic flux stimulation rescues their GSIS deficit. These findings link the β-cell functional maturation process to circadian optimization of bioenergetics, revealing a modular hierarchy by which the clock programs metabolic tissue specialization.

## Results

### Circadian clock regulator DEC1 defines mature β cells and regulates glucose homeostasis

We found that *Dec1* expression discerns mature (juvenile/adult) from immature (fetal/neonatal) β cells in mice^37^, rats^38^, pigs^39^, and humans^40^ (Fig. 1A, Extended Data Fig. 1A, Table S1). Across these species, *Dec1*’s selectivity for adult over fetal β cells is second only to *Mafa* (a key driver of mature β-cell physiology^41,42^) among adult-restricted transcription factors known to regulate human β-cell maturation. *Dec1* expression follows a 24-hour rhythm in purified adult mouse β-cells (Fig. 1B) and in primary human islets (Extended Data Fig. 1B), consistent with *Dec1*’s known role in regulating the mammalian circadian clock^13,31^. We verified that DEC1 protein is induced postnatally in mice, akin to MAFA (Extended Data Fig. 1C). To obtain initial indication of DEC1’s role in metabolism, we compared glucose homeostasis in *Dec1^-/-^* and *Dec1^+/+^* littermate mice^43^ (Fig. 1C-G). 2-3-month-old *Dec1^-/-^* mice show significantly elevated fasting and fed blood glucose (Fig. 1D), despite normal islet formation, mass, architecture, β-cell proliferation and survival (Fig. 1C, Extended Data Fig. 1D). *Dec1^-/-^* mice also display pronounced glucose intolerance at 2-3-month-old (Fig. 1E) which persists after 8 months of age (Extended Data Fig. 1E). Importantly, impaired glucose homeostasis in *Dec1^-/-^* mice is not due to obesity or insulin resistance. Body weight is indistinguishable between *Dec1^-/-^* and *Dec1^+/+^* littermates, as is body composition (Extended Data Fig. 1F, G). Insulin tolerance tests show regular sensitivity to exogenous insulin, which normalizes the elevated blood glucose observed prior to insulin injection (Fig. 1G). In contrast, *Dec1^-/-^* mice show a disrupted rise of plasma insulin during the dark (active) phase (Extended Data Fig. 1H). We traced this to significantly impaired plasma insulin/C-peptide excursions upon glucose challenge in both young (Extended Data Fig. 1I) and aged mice (Fig. 1F). These results suggest that compromised glucose homeostasis upon *Dec1* loss follows a primary defect in β-cell secretory output. Supporting this notion, we find that β-cell *DEC1* levels correlate significantly with insulin output across healthy human islet donors (but not islet autoantibody-positive, type 1, or type 2 diabetic donors), while CRISPR-Cas9-mediated *DEC1* disruption in primary human islets limits GSIS without altering insulin content (Extended Data Fig. 1J-M).

**Fig. 1.**
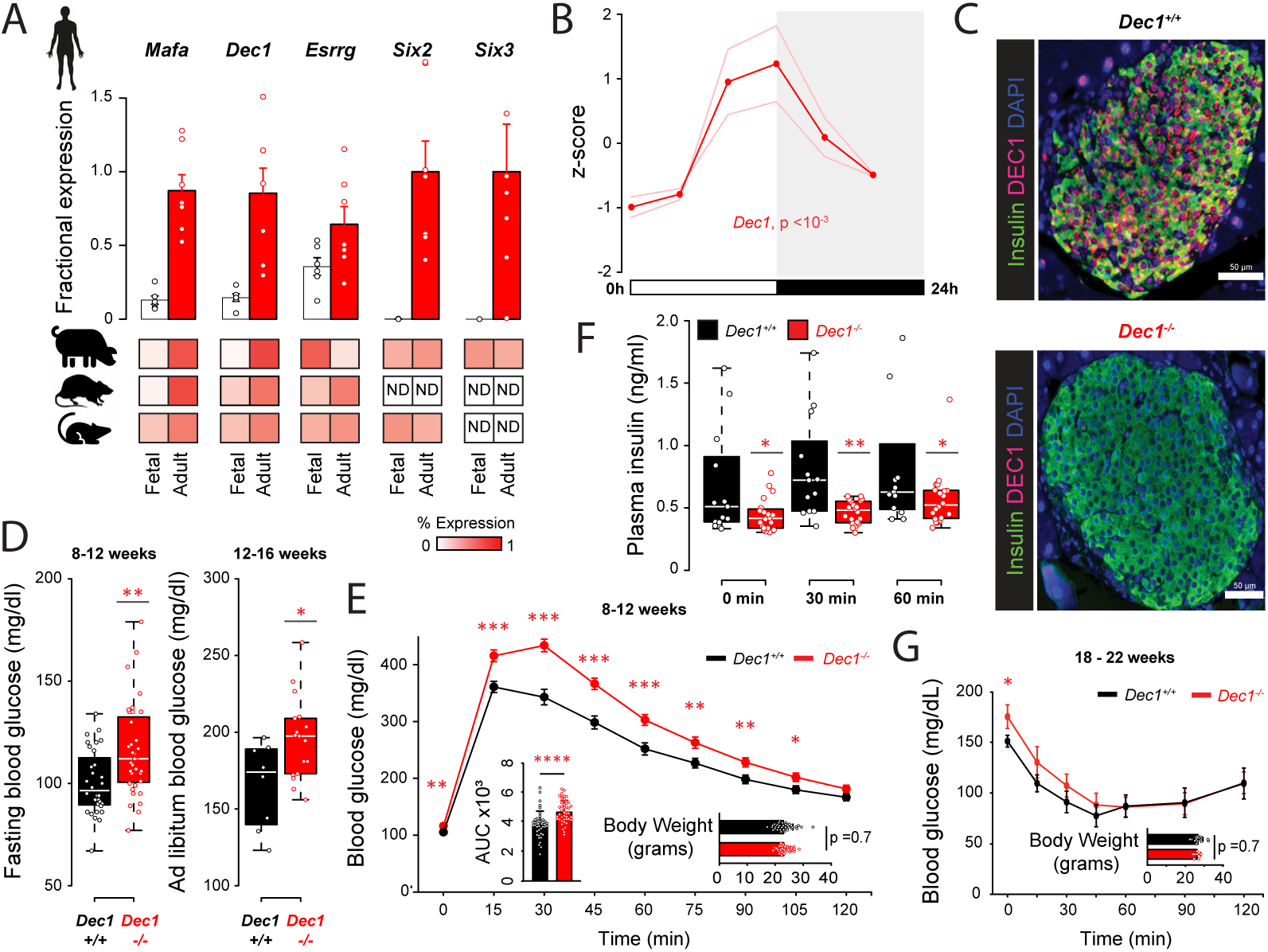
DEC1 defines mature β cells and controls blood glucose homeostasis. **(A)** Specificity of *Dec1* transcription for adult β cells compared to adult-restricted transcription factors known to regulate human β-cell functional maturation. Data are mean ±SEM fractional expression level across fetal and adult stages (see Methods) from unsynchronized β cells in human (12-18 gestational weeks, N =6; 4-60 years, N =7)^40^; pig (embryonic day 40-70, N =11; postnatal day 22, N =3)^39^; rat (embryonic day 19, N =3; postnatal week 2, N =3)^38^; and mouse (embryonic day 12.5-15.5, N =4; postnatal day 16 to 182 days, N =7)^37^. (**B**) *Dec1* oscillation in adult mouse β-cells isolated every 4h for a day^122^. Data are mean of n =2 replicate measurements, each pooling N =6 mice. Rhythmicity p-value, RAIN. (**C**) Immunofluorescence staining of insulin (green), DEC1 (red), and DAPI (blue) in pancreata from N =2 *Dec1^+/+^*and 2 *Dec1^-/-^* 8-month-old littermate mice, acquired at 20x magnification. (**D**) Blood glucose in 14-hour fasted N =32 *Dec1^+/+^*and 36 *Dec1^-/-^* 2-3-month-old littermate mice at ZT2 (left), and in *ad libitum* fed N =8 *Dec1^+/+^* and 17 *Dec1^-/-^*3-4-month-old littermate mice at ZT12 (right). (**E**) Glucose tolerance in 14-hour fasted N =39 *Dec1^+/+^* and 42 *Dec1^-/-^* 2-3-month-old littermate mice following intraperitoneal glucose administration at ZT2. Data are mean ±SEM. (**F**) Plasma insulin in 14-hour fasted N =13 *Dec1^+/+^* and 20 *Dec1^-/-^*8-month-old littermate mice following intraperitoneal glucose administration at ZT2. **(G)** Insulin tolerance in 4-hour fasted N =10 *Dec1^+/+^* and 7 *Dec1^-/-^* 4-5-month-old littermate mice following intraperitoneal insulin administration at ZT2. Data are mean ±SEM. *p <0.05, **p <0.01, ***p <0.001, ****p <0.0001, two-tailed t test [(D), (E), (G)] and Wilcoxon test (F).

Together, these findings identify DEC1 as a rhythmic transcriptional regulator selective for adult over fetal β cells and highlight its importance for physiological glucose homeostasis.

### DEC1 regulates circadian energy expenditure rhythms

To investigate energy balance, we conducted indirect calorimetry in *Dec1^-/-^* and *Dec1^+/+^*littermate mice under light-dark (LD) cycles and under dark-dark (DD) conditions that enable assessing self-sustained rhythms independent of light cues (Fig. 2A, D). We find that *Dec1^-/-^*mice keep circadian cycles but are hypo-metabolic, as they spend less energy despite increased pedestrian locomotion. In LD, analysis of energy expenditure covariance with genotype, body mass, and their interaction reveals that *Dec1^-/-^* mice expend significantly lower energy in the dark (active) phase (Fig. 2B, C). Dark-phase oxygen consumption and carbon dioxide production are also reduced, while food and water intake and the respiratory exchange ratio remain unchanged (Fig. 2A, Extended Data Fig. 2A-D, 2G). We verified that activity rhythms remain similar, with no significant differences in wheel running or overall locomotor activity (Fig. 2A, Extended Data Fig. 2H-J). Yet, pedestrian locomotion and distance traveled along the horizontal plane are increased in the light (inactive) phase (Extended Data Fig. 2E-F), suggesting increased exploration efficiency. In constant darkness (DD), *Dec1^-/-^* mice keep daily activity and energy balance rhythms but remain hypometabolic. Energy expenditure is significantly dulled in the subjective active phase and differently affected by mass (Fig. 2E, F), despite no changes in food and water intake, respiratory exchange ratio, or wheel running and overall locomotor activity levels, while ambulation remains increased throughout the day (Extended Data Fig. 2E-J).

**Fig. 2.**
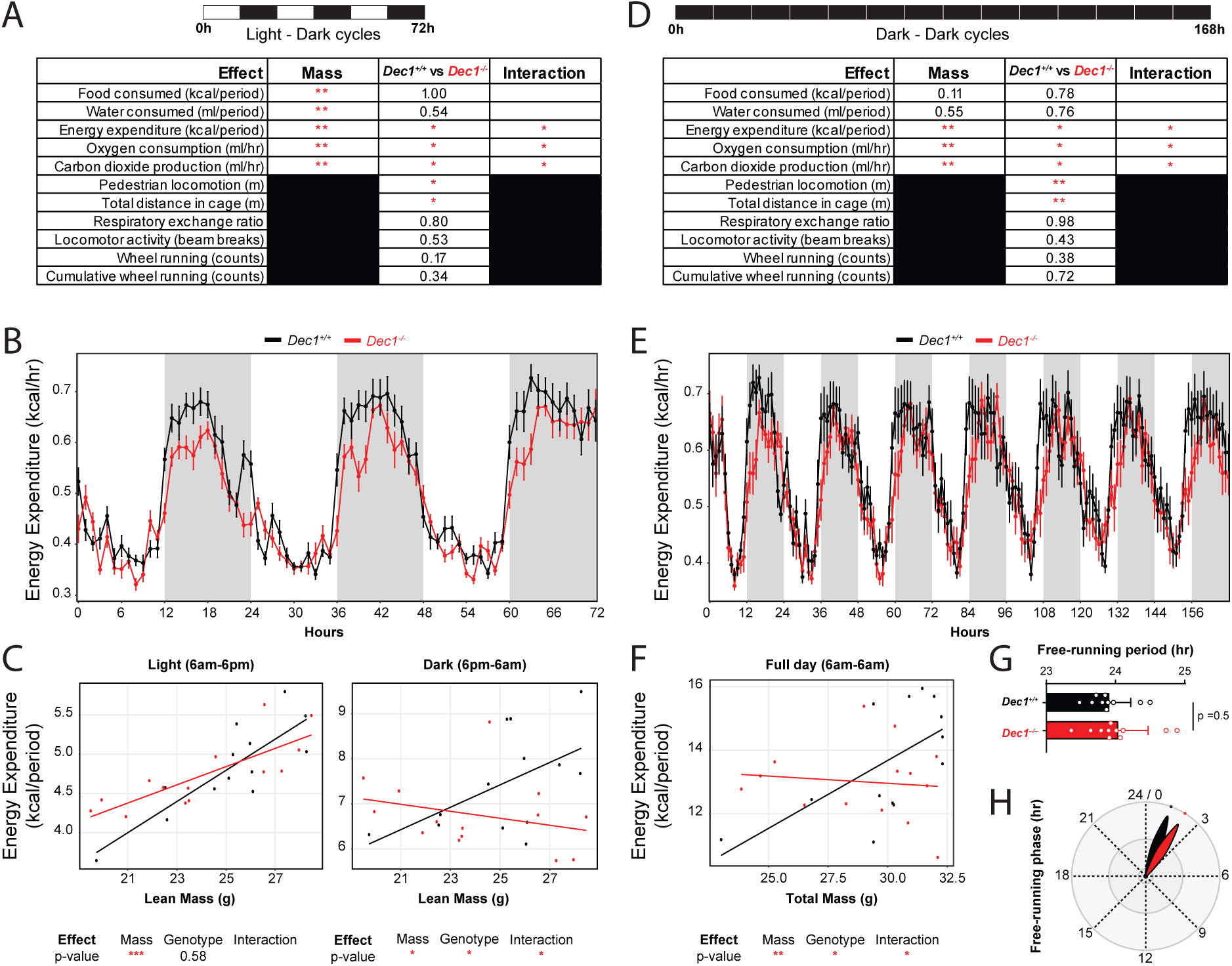
DEC1 regulates circadian energy expenditure rhythms. (**A**) Summary of results from indirect calorimetry experiments for N =13 *Dec1^+/+^* and 14 *Dec1^-/-^*8-month-old littermate mice across 3 days in 12-hour light-dark cycles. Shown are p-values corresponding to model coefficients for genotype effects, along with the mass and interaction effects for mass-dependent measurements. (**B**) Daily energy expenditure rhythms of mice from (A) across 3 days in 12-hour light-dark cycles. Data are mean ±SEM. (**C**) Regression analysis of energy expenditure and body mass by photoperiod. Model coefficients for mass, genotype, and interaction effects are indicated below. (**D**) Summary of results from indirect calorimetry experiments for N =13 *Dec1^+/+^*and 14 *Dec1^-/-^* 8-month-old littermate mice across 7 days in 12-hour dark-dark cycles (constant darkness). Shown are p-values corresponding to model coefficients for genotype effects, along with the mass and interaction effects for mass-dependent measurements. (**E**) Daily energy expenditure rhythms of mice from (D) across 7 days in 12-hour dark-dark cycles. Data are mean ±SEM. (**F**) Regression analysis of energy expenditure as a function of body mass for the full circadian cycle. Model coefficients for mass, genotype, and interaction effects are indicated below. (**G-H**) Free-running period (G) and phase (H) of wheel running rhythms of mice from (D) across 7 days in 12-hour dark-dark cycles. *p <0.05, **p <0.01, ***p <0.001, ANOVA test.

Energy expenditure rhythms show no significant change in the free-running period length (Fig. 2G), whereas the phase exhibits a delayed onset (Fig. 2H), consistent with DEC1’s known role in regulating circadian phase^32,44^.

Taken together with stable fuel preference, feeding, and overall activity (Fig. 2A, D) and unaltered body mass and composition (Extended Data Fig. 1F, G), the circadian phase-dependent hypometabolic state of *Dec1^-/-^* mice suggests inefficient energy expenditure, in line with insulin deficiency (Fig. 1F, Extended Data Fig. 2I). These findings link DEC1 to the circadian control of systemic energy metabolism.

### DEC1 selectively binds rhythmic energy metabolism and secretory genes in β cells

To better understand how DEC1 tunes β-cell function, we examined its genome-wide binding by chromatin immunoprecipitation-sequencing in unsynchronized mouse β-TC-6 cells (Fig. 3A), along with that of homolog DEC2 (also called BHLHE41 or SHARP1) (Table S2). DEC1 and DEC2 homodimerize via HLH regions that share 97% amino acid identity^45,46^, and DEC1-DEC2 heterodimers have been reported^47^, though their physiologic relevance is unknown. As expected, DEC1 and DEC2 bind CACGTG E-box motifs with significant overlap, mainly at promoters, including those of core clock activators (*Clock/Bmal1*), repressors (*Per*/*Cry*), and interlocking loop (*Ror*, *Rev-erb*, *Dec*) genes (Extended Data Fig. 3A-E). Among sites bound by DEC1 and DEC2, 26% (2,179) are also bound by CLOCK/BMAL1 in Beta-TC6 cells^29^, consistent with competition for E-box binding^31^, and enrich for circadian clock and NOTCH/TGF-β signaling effectors (Fig. 3A cluster IV, Fig. 3B, Extended Data Fig. 3F). Surprisingly, the non-overlapping DEC1/DEC2 and CLOCK/BMAL1 binding sites target largely distinct gene pathways. The DEC1/DEC2-specific cistrome enriches for (i) key maturity controllers like *Neurod1* and *Mafa*^41,48^; (ii) energy metabolism effectors like *Glut2* and *ChREBP* that enact β-cell glucose import^49^ and mitogenic signaling^50^; and (iii) components of insulin signaling (*Insr, Irs1, Igf2*) and of secretory vesicle membrane docking/fusion, including many voltage-gated K^+^/Ca^2+^ channel subunits, calmodulins/Ca^2+^-dependent kinases, and SNAREs (*Vamp2/3, Stx4a/6/16, Stxbp1/5l/6, Snap25/91/in*) (Fig. 3A clusters I-III, Extended Data Fig. 3E). By contrast, CLOCK/BMAL1-specific targets enrich for late secretory genes, including cAMP-coupled signaling mediators of vesicle budding (Fig. 3A cluster VI), consistent with impaired cAMP-mediated stimulation of insulin release in *Bmal1*-disrupted mice^29^.

**Fig. 3.**
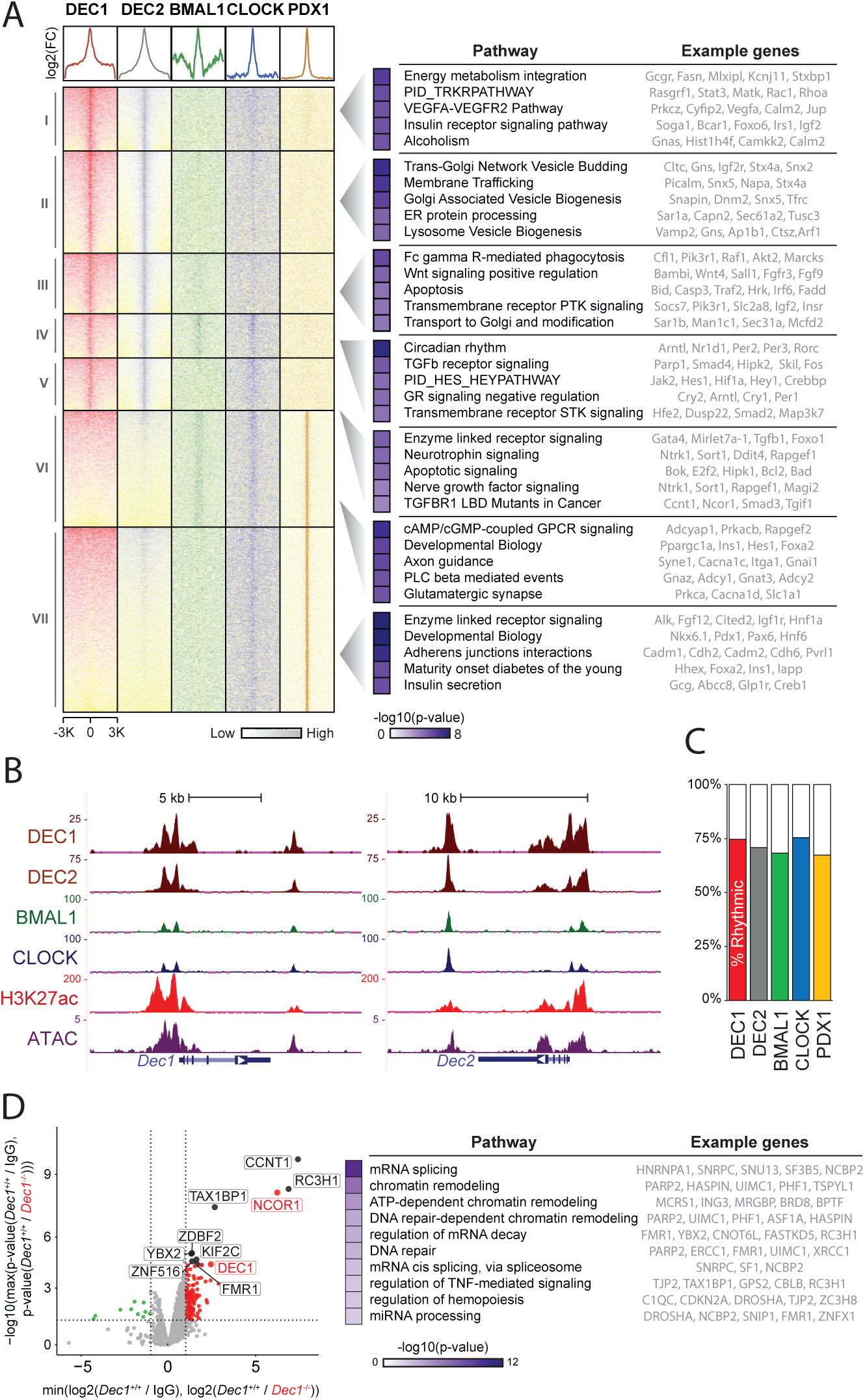
DEC1 selectively binds rhythmic energy metabolism and secretory genes in β-cells. (**A**) Left: density of ChIP-seq signal enrichment over input ±3kb around the center of each DEC1, DEC2, BMAL1, CLOCK, and PDX1 binding peak in β-cells. Roman numerals define clusters of ChIP-seq signal across the union set of binding peaks. Right: top 5 pathways enriched among genes linked to DEC1, DEC2, BMAL1, CLOCK, and PDX1 binding clusters in β-cells. (**B**) Genomic view of DEC1, DEC2, BMAL1, and CLOCK binding at *Dec1* and *Dec2* loci in β-cells. Tracks display normalized ChIP and ATAC sequencing read density. Gene models are shown below. (**C**) Fraction of DEC1, DEC2, BMAL1, CLOCK, and PDX1 target genes expressed rhythmically in β-cells. Rhythmicity analysis, dryR. (**D**) DEC1 interactors identified by IP-MS. Volcano plot shows differential enrichment of proteins detected by DEC1 IP in *Dec1*^+/+^ β-cells relative to both IgG IP in *Dec1*^+/+^ β-cells and DEC1 IP in *Dec1*^-/-^ β-cells. DEC1 and NCOR1 are labeled in red, with the ten other top enriched proteins labeled in black. Right: top 10 pathways enriched among DEC1 interactors.

DEC1/DEC2 and CLOCK/BMAL1 may cooperate with distinct co-factors to bind distinct E-boxes. For example, sites bound by CLOCK/BMAL1 but not DEC1/DEC2 are often co-bound by the pancreatic lineage-determining factor PDX1^29^ (Fig. 3A clusters VI-VII). We thus examined transcription factor binding motifs enriched across the binding clusters in Fig. 3A (Extended Data Fig. 3G). As expected, sites bound by both DEC1/DEC2 and CLOCK/BMAL1 (clusters IV-V) enrich for motifs of known circadian clock regulators (e.g., NPAS2, MYC, MYCN, CREB3L1). Whereas sites bound by DEC1/DEC2 but not CLOCK/BMAL1 (clusters I-III) enrich for motifs of E-box binding metabolic regulators (e.g., HIF1A, ChREBP, MAX). By contrast, sites bound by CLOCK/BMAL1 but not DEC1/DEC2 (clusters VI-VII) enrich for motifs of β-cell development regulators (e.g., PDX1, NKX6.1, PAX6, HNF1A/6). This suggests a basis for modular control of β-cell clock outputs, where CLOCK/BMAL1 team with pancreatic lineage-determining factors to target late secretory genes, while DEC1/DEC2 team with E-box binding metabolic regulators to target energy metabolism / signaling mediators of insulin responses to glucose.

To test rhythmicity among DEC-bound gene targets, we performed RNA-sequencing around the 24-hour clock in islets from wild-type mice and used dryR^51^ for Bayesian statistics-based rhythm model selection. DEC1/DEC2 targets show pervasive 24-hour cycling, comprising∼73% of bound and expressed genes, comparable to ∼72% for CLOCK/BMAL1 targets, which co-varied in rhythm amplitude and significance values (Fig. 3C, Extended Data Fig. 3H).

Notably, rhythmic DEC1 targets include regulators of β-cell function, energy metabolism, and insulin secretion (see below) known to oscillate in human islets^29,36,52^.

To identify DEC1-interacting proteins in β-cells, we performed DEC1 co-immunoprecipitation followed by mass spectrometry in unsynchronized mouse MIN6 cells (Fig. 3D, Extended Data Fig. 3I). As controls, we performed DEC1 purifications in *Dec1^-/-^* MIN6 cells (Extended Data Fig. 8A), and IgG purifications. We found that DEC1-specific interactors enrich for chromatin remodeling functions, including ATP-dependent remodelers like BRD8 and DNA repair-dependent remodelers like RAP80, and for mRNA processing roles, including mRNA splicing and decay factors. We validated the top-enriched chromatin-associated transcriptional corepressor (NCOR1) by co-immunoprecipitation followed by wester blotting (Extended Data Fig. 3J). Further, we found a significant overlap between DEC1’s binding peaks in β cells and those of NCOR1 in mouse liver (Extended Data Fig. 3J). This suggests that DEC1 coordinates transcriptional timing by recruiting chromatin remodelers through the NCOR1 corepressor.

Together, these findings indicate that DEC1 regulates specific steps in the GSIS pathway, by selectively binding (relative to CLOCK/BMAL1) to rhythmically expressed genes that mediate glucose and insulin metabolism, revealing a hierarchy for how distinct clock factors link to the physiology of adult β cells.

### β-cell DEC1 loss causes insulin deficiency and glucose intolerance due to islet immaturity

We next sought to ascertain the direct role of β-cell DEC1 in metabolic homeostasis by creating *InsCre Dec1^flx/flx^* (hereafter *βKO*) mice (Fig. 4A, Extended Data Fig. 4A). The mice display normal islet morphometry and architecture, insulin content, and β-cell proliferation and viability (Fig. 4A, Extended Data Fig. 4A-D). Yet, *Dec1-βKO* mice show significant glucose intolerance relative to *Dec1^flx/flx^*, *Dec1^flx/+^*, and *InsCre Dec1^flx/+^* 3-4-month-old littermates at ZT2 (Fig. 4B), which persists after 8 months of age (Extended Data Fig. 4G). While body weight and insulin sensitivity are normal (Fig. 4B, Extended Data Fig. 4H), *Dec1-βKO* mice are hypo-insulinemic, as indicated by reduced *ad libitum* plasma insulin levels and plasma insulin excursions upon glucose challenge mainly at ZT2 (Fig. 4C, Extended Data Fig. 4I-J). Further, isolated *Dec1-βKO* islets show stunted GSIS magnitude and kinetics in static and dynamic assays (Fig. 4D, Extended Data Fig. 4K). Thus, β-cell *Dec1* loss significantly hampers islet GSIS, rendering mice glucose intolerant.

**Fig. 4.**
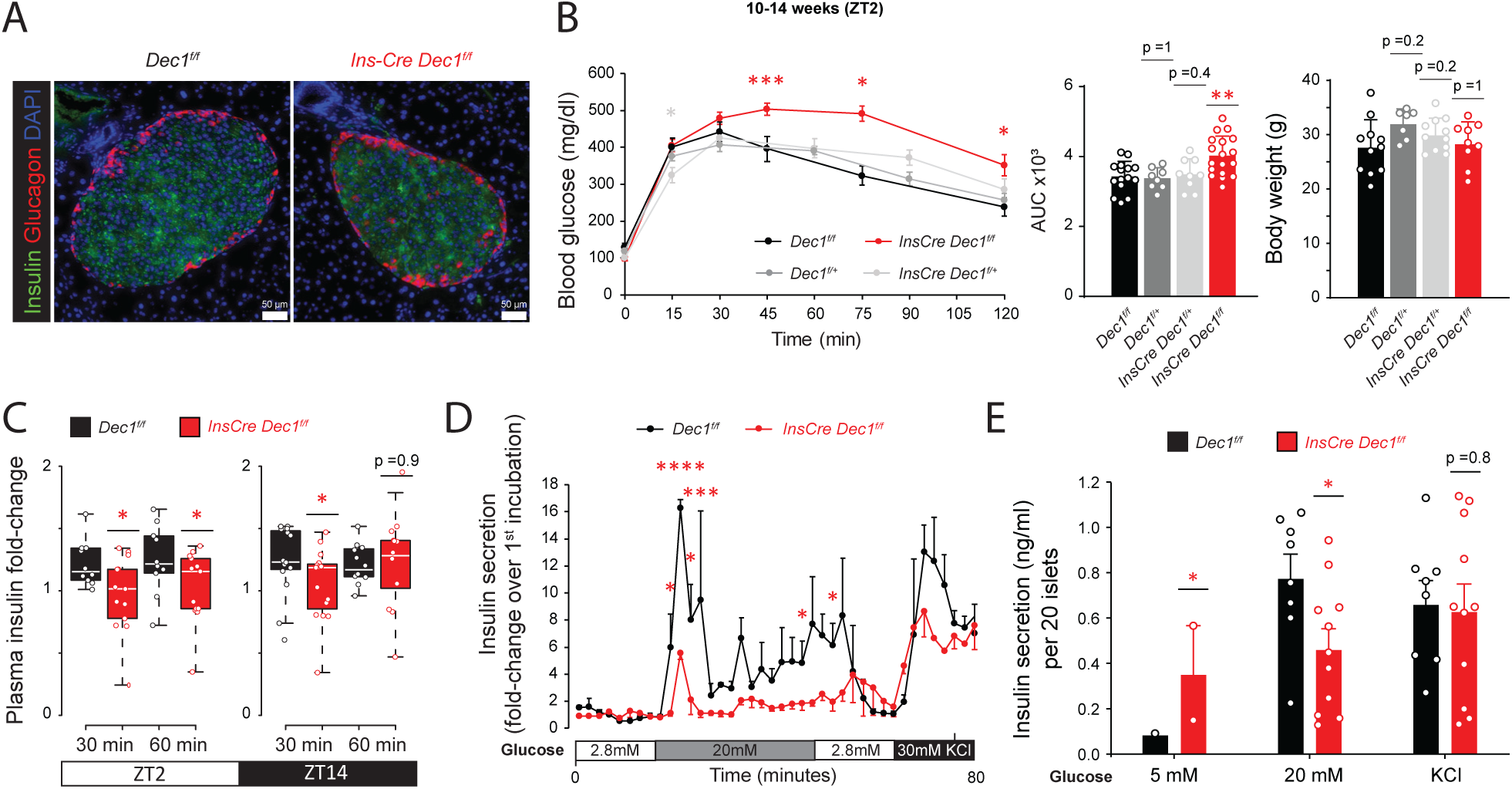
β-cell DEC1 loss causes insulin deficiency and glucose intolerance due to islet immaturity. (**A**) Immunofluorescence staining of insulin (green), glucagon (red), and DAPI (blue) in pancreata from N =2 *Dec1^f/f^* and 2 *InsCre Dec1^f/f^* 5-month-old littermate mice, acquired at 20x magnification. (**B**) Glucose tolerance in 14-hour fasted *InsCre Dec1^f/f^* (N =20), *Dec1^f/f^*(N = 17), *InsCre Dec1^f/^*^+^ (N = 12), and *Dec1^f/^*^+^ (N = 9) 3-month-old littermate mice following intraperitoneal glucose administration at ZT2. Data are mean ±SEM. (**C**) Plasma insulin fold-change in 14-hour fasted N =13-15 *InsCre Dec1^f/f^* and 10-15 *Dec1^f/f^* 2-3-month-old littermate mice following intraperitoneal glucose administration at ZT2 (left) and at ZT14 (right). (**D**) Glucose-stimulated insulin secretion dynamics in perifused islets from N =2 *InsCre Dec1^f/f^* and 2 *Dec1^f/f^* 8-month-old littermate mice, each with n =2-4 replicate measurements of 20 similar-sized islets. Data are mean ±SEM, normalized to the mean of the first incubation. (**E**) Insulin secretion of islets isolated from N =10 *InsCre Dec1^f/f^* and 9 *Dec1^f/f^* 8-10-month-old littermate mice, each with n =2 replicate measurements of 20 size-matched islets under the indicated parallel static incubations. Data are mean ±SEM. *p <0.05, **p <0.01, ***p <0.001, ****p <0.0001, two-tailed t test

Islets develop mature GSIS responsiveness after birth, marked by increases in the glucose threshold for insulin release and the glucose-coupled release capacity^21,22^. To define the defective step in *Dec1-βKO* islet GSIS, we tested their responses across increasing glucose levels in static and dynamic incubations. Isolated *Dec1-βKO* islets show elevated insulin secretion under sub-physiological glucose, mounting anomalous insulin responses to 2.8 and 5mM glucose (Fig. 4E, Extended Data Fig. 4K). This indicates a lower glucose threshold for insulin release, reminiscent of fetal/neonatal islets^21,22^. By contrast, insulin responses to saturating (20mM) glucose levels are significantly lower in *Dec1-βKO* islets, despite normal insulin content and release upon chemical depolarization with KCl (Fig. 4D-E, Extended Data Fig. 4K-L). This indicates blunted glucose-coupled insulin release capacity. We verified that the glucose threshold for insulin release and the release capacity are also diminished upon CRISPR-Cas9-mediated *DEC1* disruption in primary human islets (Extended Data Fig. 1J, L).

Collectively, these findings reveal that β-cell DEC1 loss results in insulin deficiency owing to functionally immature glucose responsiveness of pancreatic islets.

### DEC1 synchronizes metabolic and insulin secretion gene rhythms in β cells

To assess how DEC1 regulates mature GSIS function, we examined the impact of β-cell *Dec1* loss on islet gene expression patterns. RNA-sequencing over a 24-hour cycle in islets isolated and pooled from *Dec1-βKO* mice revealed 3,176 genes (13% of all detected) with differential rhythms (Fig. 5A, Extended Data Fig. 5A, Table S3). Rhythm-disrupted genes enrich for functions in vesicular transport and epigenetic regulation (Fig. 5A). These genes include *Bmal1* (Fig. 5E), although other core clock genes that are also bound by DEC1 lack significant rhythm changes (Extended Data Fig. 5F). We also found 818 genes that gain a 24-hour rhythm, enriching for calcium signaling mediators and general machinery for vesicle budding. Cycling amplitudes of rhythm-disrupted and-gained genes are globally greater than those of rhythm-retaining genes (Extended Data Fig. 5A), supporting their *bona-fide* rhythmicity. Intriguingly, many GSIS regulators retained daily cycling but became phase shifted (Fig. 5A-C, Extended Data Fig. 5C-E). These include (i) effectors of glucose sensing (*Glut2*, *Foxo1*) and mitochondrial metabolism (TCA, electron transport chain, ATP synthesis enzymes); (ii) PI3K-Akt-mTOR signaling modulators of energy homeostasis (*Gsk3b, Igf1*); and (iii) direct effectors of insulin transport (tubulins, ARF/RAB GTPases) and exocytosis (G-protein subunits, *Irs1/2, Syt4/13*). Disparate phase shifts include phase advances for electron transport chain components and phase delays for the *Glut2*/*Igf1* effectors of glucose sensing (Fig. 5B-C, Extended Data Fig. 5D, E, G). Globally, phase offsets between respiratory metabolism and GSIS gene rhythms are lost in *Dec1-βKO* islets (Extended Data Fig. 5D), indicating time desegregation of metabolic processes.

**Fig. 5.**
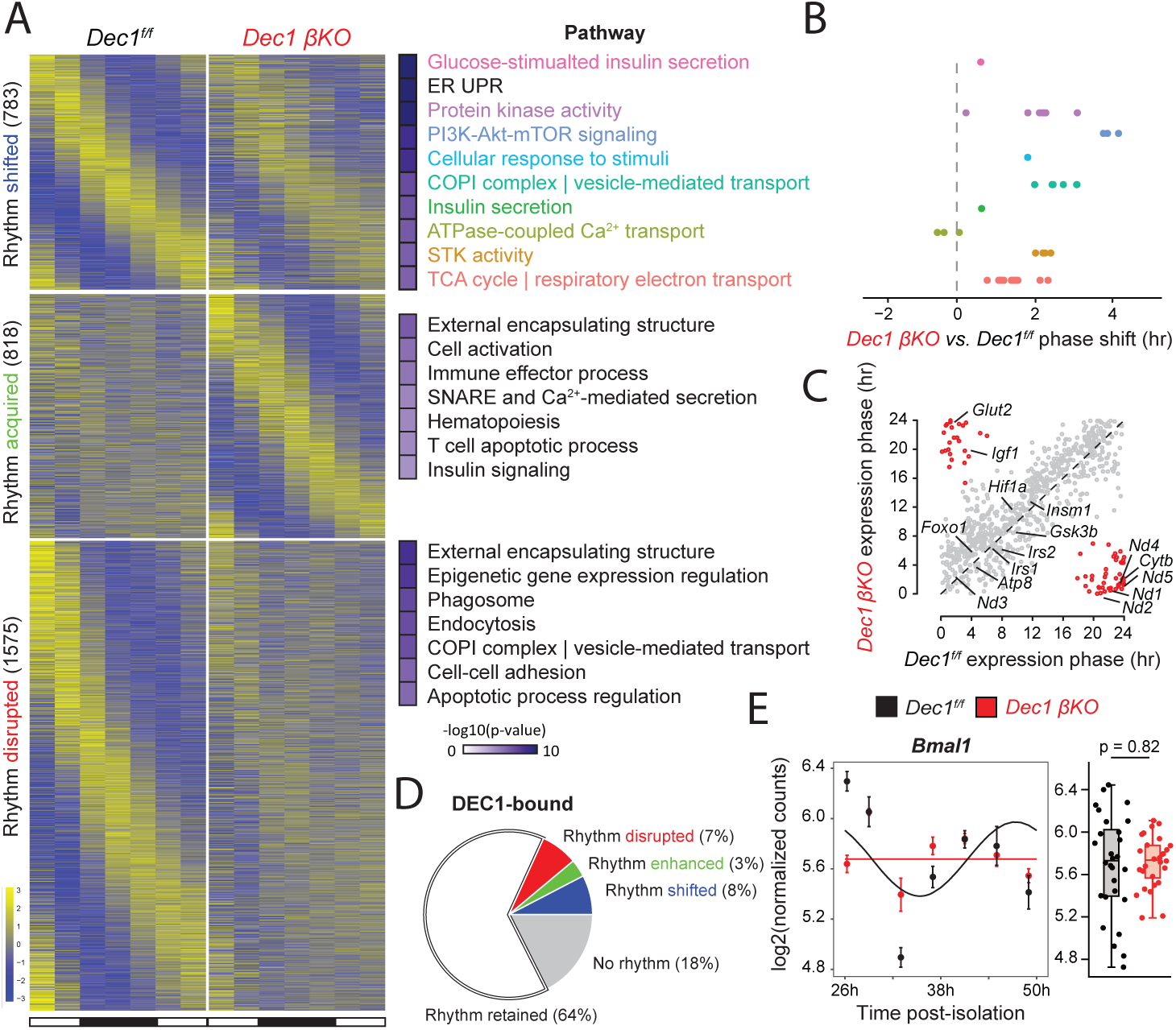
DEC1 synchronizes respiratory metabolism and insulin secretion gene rhythms in pancreatic β-cells. (**A-B**) Expression of genes whose rhythm is disrupted (bottom), enhanced (middle), or shifted (top) in islets isolated and pooled from N =2 *Dec1*-*βKO* and 2 *Dec1^f/f^* 7-month-old littermate mice (A, left), highlighting enriched pathways (A, right), and relative phase change for pathways enriched among rhythm-shifted genes, colored by biological theme (B), with each dot representing the average relative phase change among the genes driving enrichment for specific pathways within each biological theme. Data are mean of n =4 replicate measurements. (**C**) Phase of expression, defined as the time of peak expression, for the 783 rhythm-shifted genes from (A). Highlighted in red are 70 genes with a >12hr difference in expression phase, which include mediators of glucose sensing, metabolism, and insulin secretion. (**D**) Among DEC1-bound genes, the fraction showing non-rhythmic expression (gray) or rhythmic expression that is retained (white), disrupted (red), enhanced (green), or phase-shifted (blue) in *Dec1*-*βKO* islets from (A) is shown. (**E**) *Bmal1* temporal expression pattern in *Dec1*-*βKO* islets from (A). Data are mean ±SEM from n =4 replicate measurements. p-value, two-tailed t test.

Genes with differential rhythms that are also bound by DEC1 are enriched for regulatory roles in glucose responsiveness and insulin secretion, driven by genes like *Hif1a*, *Irs1/2*, *Mafa*, *Pdk1*, and *Pkm* (Extended Data Fig. 5B). Among genes with differential rhythms, rhythm-shifted genes are overrepresented for DEC1 binding (Extended Data Fig. 5B), consistent with its known role in coordinating the oscillation phase of its targets^44^.

Together, these findings indicate that synchronizing insulin responses with glucose sensing and oxidative metabolism is an important temporal gene-control function of DEC1 in mature β cells.

### DEC1 regulates coupling of insulin secretion to glucose, shaping islet cell-cell coordination

Islets rely on temporally orchestrated interactions among heterogeneous cell types to sustain glucose homeostasis^53,54^. To better understand cell type-specific effects in *Dec1-βKO* islets, we subjected them to single-cell RNA sequencing (scRNA-seq), single-cell assay for transposase-accessible chromatin by sequencing (scATAC-seq), and single-cell proteomic mass spectrometry (SCoPE-MS)^55,56^. Unsupervised clustering followed by marker-based cross-modal integration^57^ identified endocrine (α, β, δ, γ), vascular, stellate, and immune cells (Fig. 6A, Extended Data Fig. 6A, B). These included as endocrine subpopulations (i) β1/β2 cells, which show higher/lower expression of insulin and the maturity markers *Ucn3*^22^ and *Iapp*^58^; (ii) α1/α2 cells, which show higher/lower glucagon and *Irx2, Arx*; (iii) δ1/δ2 cells, which show higher/lower somatostatin and *Hhex*. We verified that *Dec1* is enriched in β cells relative to other endocrine cells and expressed at levels comparable to brain neurons and immune cells, where its function has been studied (Extended Data Fig. 6E).

**Fig. 6.**
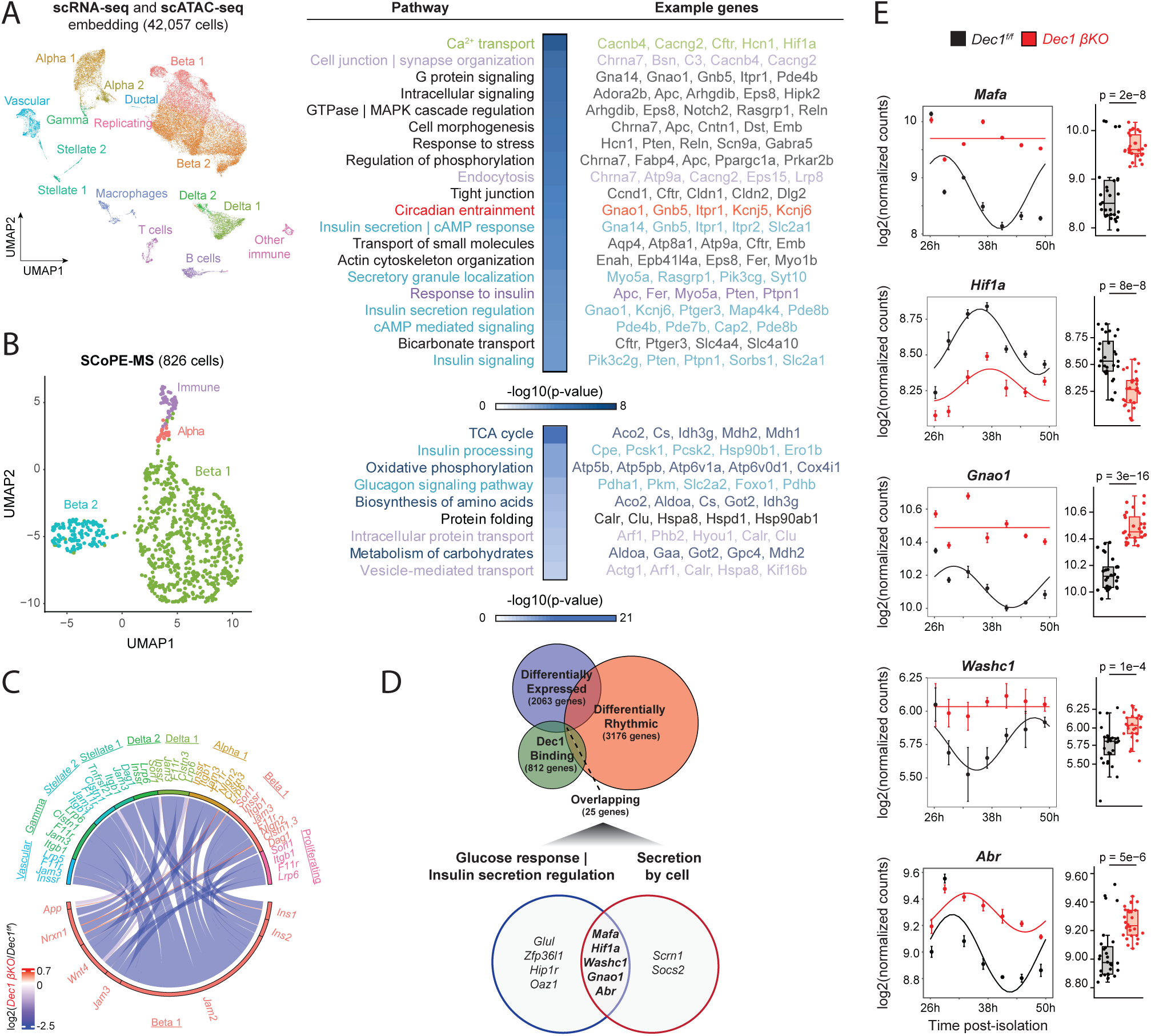
DEC1 regulates coupling of insulin secretion to glucose, shaping islet cell-cell coordination. (**A**) Co-embedding of single-cell open chromatin (scATAC-seq) and gene expression (scRNA-seq) profiles in islets isolated from N =2 *Dec1*-*βKO* and 2 *Dec1^f/f^* 7-month-old littermate mice, each sequenced separately (left). Pathways enriched among genes showing both differential expression and differential chromatin accessibility (right). (**B**) Single-cell clustering by protein expression profiles, assayed by SCoPE-MS, in islets isolated from N =2 *Dec1*-*βKO* and 2 *Dec1^f/f^* 8-month-old littermate mice, each run separately (left). Pathways enriched among differentially expressed proteins (right). (**C**) Ligand-receptor interaction analysis (see Methods) showing 7 of the top 10 differentially active interactions, where ligand expression is significantly altered in Beta 1 cells from (A) with higher expression of insulin and maturity markers. (**D**) Overlap between DEC1-bound genes in β cells and genes that are differentially rhythmic and differentially expressed in β cells from *Dec1*-*βKO* mice, highlighting the top 2 pathways enriched among the 25 genes shared by all three datasets, which include genes mediating glucose responsiveness and insulin secretion. (**E**) Temporal expression pattern of *Mafa, Hif1a, Gnao1, Washc1,* and *Abr* in islets isolated and pooled from N =2 *Dec1*-*βKO* and 2 *Dec1^f/f^* 7-month-old littermate mice from Fig. 5. Data are mean ±SEM from n =4 replicate measurements. p-value, two-tailed t test.

We find that β1 cells are significantly depleted in *Dec1-βKO* islets (Extended Data Fig. 6A), while overall endocrine cell type ratios remain intact, indicating selective loss of β cells in functionally mature states. In line with this notion, *DEC1* disruption downregulates key β-cell maturity genes, including *MAFA* and *SIX2*, in primary human islets (Extended Data Fig. 1K, L). We also verified blunted postnatal induction and overall downregulation of β-cell MAFA protein in pancreata from our mouse *Dec1* knockout models (Extended Data Fig. 1C, 4E).

Analyzing RNA changes across *Dec1-βKO* islet cell groups reveals pervasive gene deregulation, with ∼40% of changes equally ascribed to β1 and β2 cells (Extended Data Fig. 6C). Genes showing both differential expression and differential chromatin accessibility in β cells are enriched for functions in GSIS regulation, including Ca^2+^ transport, cAMP signaling, secretory granule localization, and insulin secretion (Fig. 6A, Extended Data Fig. 6D). Critically, these changes remodel the proteome, as single-cell proteomics unveils significantly altered levels of proteins mediating glucose metabolism (TCA, OXPHOS enzymes) and GSIS (insulin processing, transport, and secretion) in β cells (Fig. 6B, Extended Data Fig. 6G-H, Table S4). To identify subsequent effects on intercellular signaling, we focused on probabilistic ligand-receptor interaction changes where the ligand is altered in β1 cells, given their marked downregulation (Fig. 6C, Extended Data Fig. 6F). As expected, we find reduced insulin signaling from β1 cells (to themselves and to δ cells). This is accompanied by reduced signaling of the WNT4 maturity modulator^59^, the JAM2/3 junction adhesion proteins, and the NRNX1 secretory granule co-regulator from β1 cells to other endocrine cells, while ligand activity of the APP amyloid beta precursor protein is enhanced. These results offer a molecular basis for how GSIS deregulation in β cells disrupts intercellular coordination in *Dec1-βKO* islets.

We next studied the impact of altered gene expression on rhythmicity in *Dec1-βKO* islets. Most differentially expressed genes either lack or retain rhythmicity (Extended Data Fig. 6I). Yet, specific subsets, enriched for GSIS effectors, change in both rhythm and mean expression level, like exocytic machinery and genes involved in fatty acid and tryptophan metabolism, which are both upregulated and phase-shifted in rhythm (Extended Data Fig. 6J). To establish direct transcriptional regulation by DEC1, we intersected differentially expressed and rhythm-altered genes with the DEC1 cistrome (Table S5). This identified 25 DEC1 targets enriched for roles in glucose responsiveness and insulin secretion (Fig. 6D, Table S5). Remarkably, 5 such targets drive enrichment in both the glucose-responsiveness and secretory annotation clusters (Fig. 6D, E): (i) the key β-cell maturity transcription factors *Mafa*^41,42^ and *Hif1a*^60^; (ii) the exocyst co-factor *Wash1c*^61^; and (iii) the G_o_-type G protein–coupled receptor subunit *Gnao1* and the *Abr* RhoGEF and GTPase activator.

Together, these findings reveal direct DEC1 targets influencing glucose responsiveness and insulin secretion, delineating a likely pathway for DEC1’s control of adult β-cell GSIS function.

### A deficit in circadian bioenergetics underlies immature β-cell function upon DEC1 loss

Given DEC1’s binding and rhythmic regulation of energy metabolism genes, we next tested its influence on cellular energetics. We find that DEC1 is vital for time-of-day variation in GSIS reponsiveness^29,62^ in both *Dec1*-knockout mouse islets and *DEC1*-knockdown human primary islets (Fig. 7A, Extended Data Fig. 8G). This follows DEC1’s critical requirement for circadian mitochondrial energetics. Mitochondrial fusion into tubular networks enabling higher oxidative activity is under circadian control^63^, and day-to-night switching from fragmented (spherical) to tubular (branched) mitochondria is blunted in *Dec1*-ablated β cells, despite normal mitochondrial content (Fig. 7B, Extended Data Fig. 7A, B). At 36h post-isolation, wild-type islets exhibit a >4-fold rise in GSIS responsiveness, and β cells show reduced mitochondrial sphericity and greater branching, surface area, and volume—changes not observed in *Dec1*-ablated counterparts.

**Fig. 7.**
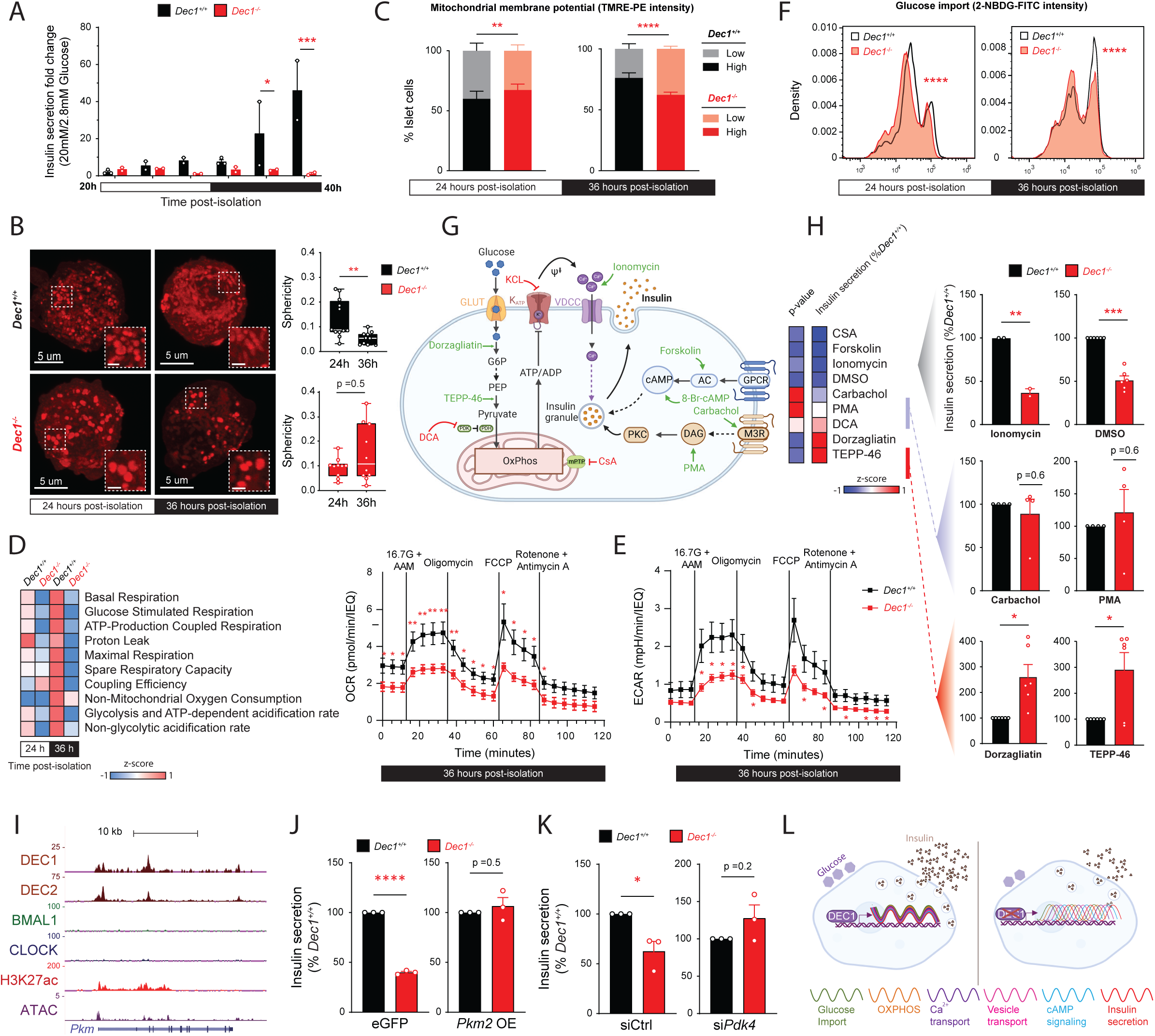
A deficit in circadian bioenergetics underlies immature β-cell function upon DEC1 loss. (**A**) Insulin secretion fold-change between 2.8mM and 20mM glucose sequential static incubations of islets isolated and pooled from N =4 *Dec1^+/+^* and 4 *Dec1^-/-^* 2-3-month-old littermate mice assayed at 4h intervals for 20h. Data are mean ±SEM from n =2-3 replicate measurements of 20 size-matched islets per time point. (**B**) Representative confocal images of mitochondria stained with PKmito Orange FX in dissociated insulin-labeled β cells from islets isolated from N =3 *Dec1^+/+^* and 2 *Dec1^-/-^*2 month-old littermate mice at 24h and 36h post-isolation, acquired at 63x magnification. Inset scale bar = 1um. Quantification of mitochondrial sphericity for each genotype and time point (right). Data are distributions of measurements across n =10-12 independent fields of view, each with >50 mitochondria. (**C**) Proportion of islet-cells isolated from N =4 *Dec1^+/+^* and 4 *Dec1^-/-^* 2-4-month-old littermate mice with low and high mitochondrial membrane potential determined by TMRE staining at 24h and 36h post-isolation. Data are mean ±SEM from n =4 replicate measurements, each with >5000 live singlet cells. (**D-E**) Metabolic flux at 24h and 36h post-isolation in islets isolated from N =5 *Dec1^+/+^* and 5 *Dec1^-/-^* 2-4-month-old littermate mice (D, left) quantified from measurements of oxygen consumption rate (OCR) (D, right) and extracellular acidification rate (ECAR) (E) in response to sequential incubations with 16.7mM glucose and a physiological amino acid mixture (Table S6) (16.7G + AAM^119^), oligomycin, FCCP, and rotenone and antimycin A at the indicated times. Data are mean ±SEM from n =4-7 replicate measurements of 12 similar-sized islets. (**F**) Glucose uptake in islet-cells isolated from N =2 *Dec1^+/+^* and 2 *Dec1^-/-^* 2-3-month-old littermate mice quantified by import of the fluorescent glucose analog 2-NBDG at 24h and 36h post-isolation. n=2 replicate measurements, each with >5000 live singlet cells. (**G-H**) Screen of insulinotropic agents to rescue the GSIS defect of isolated *Dec1*-knockout islets, created with BioRender.com (G), and results for significance and insulin secretion fold-change (20mM / 2.8mM glucose) in sequential static incubations of *Dec1^-/-^* relative to *Dec1^+/+^* islets (H, left), with the relative vehicle (DMSO) or secretagogue-stimulated insulin secretion shown to the right (H, right). Data are mean ±SEM from 2-4-month-old littermate mice (N =2-4 for each genotype), each with n =2-3 replicate measurements of size-matched islets, normalized to intracellular insulin content. (**I**) Genomic view of DEC1, DEC2, BMAL1, and CLOCK binding at the *Pkm* locus in β-cells. Tracks display normalized ChIP and ATAC sequencing read density. Gene model is shown below. (**J**) Insulin secretion fold-change between 2.8mM and 20mM glucose of islets isolated and pooled from N =6 *Dec1^+/+^* and 5 *Dec1^−/−^* 2-3-month-old littermate mice infected with lentiviral *Pkm2* overexpression plasmid or eGFP control plasmid. Data are mean ±SEM from n =3 replicate measurements. **(K)** Insulin secretion fold-change between 2.8mM and 20mM glucose of islets isolated and pooled from N =3 *Dec1^+/+^*and 3 *Dec1^−/−^* 2-3-month-old littermate mice transfected with siRNA targeting *Pdk4* (*siPdk4*) or control siRNA (siCtrl). Data are mean ±SEM from n =3 replicate measurements. (**L**) Model for DEC1-mediated coordination of mature insulin responses to glucose in β-cells via synchronization of energy metabolism, signaling, and secretory gene rhythms, which become misaligned in DEC1’s absence. *p <0.05, **p <0.01, ***p <0.001, ****p <0.0001, two-way ANOVA test [(A), (D), (E)], Pearson’s Chi-squared test [(C), (F)], two-tailed t test [(B), (H), (J), (K)].

To investigate the basis for disrupted mitochondrial dynamics in *Dec1*-ablated β cells, we generated a *Dec1^-/-^* MIN6 cell line (Extended Data Fig. 8A) that recapitulates key physiological features and facilitates biochemical studies. We used CRISPR-Cas9 gene editing and homology-directed repair to establish *Dec1* nullizygous MIN6 cells, verifying DEC1 protein loss. Whereas control MIN6 cells show a robust circadian rhythm in GSIS responsiveness, *Dec1^-/-^* MIN6 cells show no significant rhythm and dulled GSIS, reflecting elevated basal insulin release and blunted stimulated release capacity (Extended Data Fig. 8B). Using this system, we examined circadian regulation of mitochondrial fusion–fission machinery (Extended Data Fig. C-F). DRP1 phosphorylated at Ser616, marking the active, fission-promoting form of the protein shows a robust circadian rhythm in control cells, antiphasic with GSIS responsiveness (Extended Data Fig. 8C, D). In contrast, *Dec1^-/-^* cells show no significant rhythm and an overall reduction in phospho-Ser616 DRP1 levels. In parallel, the mitochondrial fusion regulators *Opa1* and *Mfn1* display rhythmic expression in control cells, in phase with GSIS responsiveness, and DEC1 loss attenuates both their rhythmicity and overall level, whereas *Mfn2* neither oscillates in control cells nor is altered by DEC1 loss (Extended Data Fig. 8F).

To assess if altered mitochondrial dynamics reflect turnover changes in *Dec1^-/-^* cells, we quantified mitophagy via PINK1–PARKIN colocalization analysis in mouse islet β cells and examined mitochondrial ultrastructure via transmission electron microscopy (TEM) in MIN6 cells (Extended Data Fig. 8H–I). Neither approach revealed significant differences, while TEM verified cycling from spherical to tubular mitochondria, in phase with GSIS responsiveness, in control but not *Dec1^-/-^* MIN6 cells. These findings indicate that altered mitochondrial dynamics in DEC1-deficient β cells follow a primary defect in circadian fusion-fission cycles rather than altered mitophagic flux.

In line with stunted mitochondrial dynamics, circadian cycling of cellular reactive oxygen species dulls in DEC1’s absence (Extended Data Fig. 7C). This coincides with sharp declines in the rise in cells with high mitochondrial membrane potential (MMP) and in maximal MMP in phase with peak GSIS responsiveness (Fig. 7C, Extended Data Fig. 7D). Measuring oxygen consumption further reveals that *Dec1* loss significantly stifles basal, glucose-stimulated, ATP-linked, and maximal respiration, hindering spare respiratory capacity, mainly in the peak GSIS phase, whereas oxidative responses to amino acids are unaltered (Fig. 7D, Extended Data Fig. 7E, H). Thus, DEC1 regulates circadian rhythms in both mitochondrial dynamics and activity and GSIS responsiveness.

Reduced mitochondrial dynamics and activity in DEC1’s absence may reflect adaptations to reduced glucose utilization. We find that extracellular acidification rates, reflecting glycolytic activation and the associated rise in ATP-dependent proton efflux, are indeed markedly stifled (Figs. 7D, E, and Extended Data Fig. 7F, I). We thus tested if curbed reliance on glucose upon *Dec1* loss reflects lower glucose import. Cells with high uptake of a fluorescent glucose analog markedly rise in the peak GSIS phase, which becomes significantly disrupted in DEC1’s absence (Fig. 7F). Taken together with compromised acidification, mitochondrial dynamics, and oxidative respiration, these findings indicate that immature GSIS upon DEC1 loss stems from a deficit in circadian glucose utilization.

We next conducted a targeted screen of insulinotropic agents to rescue the GSIS defect of isolated *Dec1^-/-^* and *Dec1-βKO* islets (Fig. 7G-H, Extended Data Fig. 7J). Glucose stimulation along with cAMP agonists forskolin or 8-br-cAMP fails to rescue the significant GSIS deficit, as does co-stimulation with ionomycin, a Ca^2+^ ionophore, in line with normal Ca^2+^ influx dynamics (Extended Data Fig. 7G). Conversely, glucose-independent stimulation with PMA (phorbol 12-myristate 13-acetate), a direct PKC activator, or with carbachol, a muscarinic receptor agonist, restores insulin secretion to wild-type levels (Fig. 7H, Extended Data Fig. 7J). To rescue the specific defect in glucose coupling of insulin release, we then sought to increase metabolic flux. Raising the pyruvate pool with TEPP-46, a pyruvate kinase (PKM) activator, or the acetyl-CoA pool with dichloroacetate, a pyruvate dehydrogenase kinase (PDK) inhibitor, dramatically boosts GSIS responsiveness beyond wild-type activity (Fig. 7H). Similarly, enhancing glycolytic flux with dorzagliatin, a glucokinase activator, improves GSIS responses beyond wild-type levels. As genetic alternatives to these pharmacologic manipulations of metabolism, we overexpressed *Pkm2* (a direct DEC1 target) and depleted *Pdk4* in *Dec1*-deficient islets (Extended Data Fig. 8I-K). In both cases, GSIS responsiveness was restored to the level of controls (Fig. 8J, K). Thus, greater energy flux relieves the hypo-metabolic state caused by *Dec1* loss, revealing a molecular basis for DEC1’s impact on β-cell maturation and systemic glucose homeostasis (Fig. 8L).

## Discussion

We have defined a genomic basis for the coordination of metabolism by adult β cell-specific circadian clockwork. Tissue-intrinsic physiology rhythms are thought to be programmed by the ubiquitous CLOCK/BMAL1 activators teaming with lineage-restricted transcription factors^4,64^, in line with the phenotypes of conditional CLOCK/BMAL1 knockouts. Yet, not all rhythmic genes are CLOCK/BMAL1 targets, and such knockouts may lack the entire circadian system^6^. Using pancreatic β cells as a model for cell type-specific circadian control of metabolism, we identify separable roles for CLOCK/BMAL1 and the adult-restricted DEC1 regulator in programming energy metabolism and secretory gene oscillations that set daily variation in the glucose responsiveness of insulin secretion.

Our genomic binding studies illuminate how β cell-specific clock outputs are generated. Both CLOCK/BMAL1 and DEC1/DEC2 target E-box motifs and bind clock genes, including each other’s, but most of their binding targets are distinct. CLOCK/BMAL1 preferentially binds sites co-targeted by pancreatic lineage-determining factors like PDX1^29^, enriched for late secretory genes. By contrast, DEC1/DEC2 preferentially binds metabolic/signaling mediators of insulin responses to glucose, co-targeted by energy metabolism regulators like HIF1A and ChREBP. As both sets of target genes show robust rhythms, our results reveal a hierarchy for coordinating clock outputs, with cell state-specific gene rhythms set mainly by interlocking loop regulators, rather than core clock components directly^65,66^. This model does not exclude indirect effects, as core clock and interlocking loop controllers regulate each other. Further studies will be needed to determine the basis for these factors targeting distinct E-boxes within the same cell type or across diverse tissues^67,68^.

Our discovery that *Dec1* loss causes lifelong glucose intolerance and insulin deficiency, as with pancreas/β-cell *Bmal1* loss but without stunted islet growth or viability^17,69^, argues that circadian GSIS control operates mainly through this arm of the clock. We find that β-cell DEC1 is activated after birth and before weaning, akin to the maturity controller MAFA. This coincides with the rise in the glucose threshold for β-cell insulin release between birth and weaning, before glucose-coupled release capacity expands upon weaning^21–23^. We further find that *Dec1* loss disrupts this progression, resulting in fully formed islets with a diminished threshold for insulin release and release capacity, reminiscent of fetal/neonatal islets^21,22^. As DEC1 regulates MAFA developmental induction, overall level, and transcription oscillation by binding the *Mafa* gene, our findings argue that DEC1 coordinates circadian control of maturation regulators to foster metabolic specialization, linking the circadian system to development of the mature β-cell phenotype.

Temporal segregation of incompatible metabolic processes is key for optimized cell and tissue physiology^1,2^. Indeed, we show that DEC1-mediated synchrony of energy metabolism rhythms is vital to coordinate the specialized physiology of β cells, consistent with DEC1’s ability to entrain circadian rhythms to environmental stimuli^13,44^. We find that circadian rhythms in energy balance and activity show delayed phase onset in *Dec1^-/-^* mice. Mechanistically, we find phase desynchrony in the islet expression rhythms of oxidative metabolism and insulin secretion effectors, resulting in circadian deficits in glucose utilization, mitochondrial activity, and GSIS responsiveness. We further find that DEC1-mediated control of mitochondrial dynamics, which are known to be circadian-regulated^63^, is key for optimizing β-cell metabolic function. DEC1-dependent transcriptional programs thus coordinate metabolic flux shifts governing β-cell function, and targeted activation of downstream pathways, such as pyruvate kinase activation, can restore functional deficits by compensating for blunted metabolic flux.

This raises the possibility of timed modulation of DEC1’s targets as a strategy to mimic DEC1’s maturational effects by stimulating circadian metabolic control in β-cells. Such a strategy may be relevant in diabetic patients, where the lost correlation between DEC1 expression and islet insulin release output suggests loss of mature GSIS responsiveness in β cells.

Optimizing tissue functions to energy availability offers a clear fitness advantage^1,2^. As circadian clocks emerge in metabolic tissues during the onset of daily feeding-fasting cycles^26,70^, an intriguing possibility is that they unlock phenotypic maturity by optimizing specialized tasks to daily energy availablity^9^. Lapses in such circadian optimization, as occurs with chronic circadian rhythm disruption^5,18^, may thus lead to multi-tissue dysfunction due to widespread loss of mature cell phenotypes.

Overall, our finding that insulin secretion’s responsiveness to glucose relies on DEC1-driven synchrony of circadian metabolic and secretory rhythms illuminates how adult β-cells achieve specialized function. Given major interest in achieving maturation of *in vitro*-derived human islets for diabetes modeling and therapy^71,72^, the fact that DEC1 marks mature β-cells from rodents to humans heralds new paths to co-opt bioenergetic rhythms to realize fully functional lab-grown β-cells for research and medicine.

### Limitations of study

The experimental systems used in this study have limitations to consider. All *in vivo* experiments were performed in male mice to minimize variability in glucose tolerance tied to estrous cycles, so sex-specific effects were not evaluated. Some of our experiments utilize human cadaveric islets, which display an inherent degree of functional variability between donors, due to individual variability in age, gender, and metabolic health. To mitigate this, we used only islets from healthy donors with a BMI within the normal range. Experiments were replicated with islets from multiple donors to ensure that findings were not donor specific. Mouse β-cell lines were used for transcription factor ChIP-sequencing, co-immunoprecipitation followed by mass spectrometry, and circadian western blotting experiments, given the substantial cell material requirements. For experiments involving β-cell *Dec1* knockout during mouse embryogenesis, we used a rat insulin promoter-driven Cre recombinase (RIP-Cre)^73^ that has been found to be expressed in the brain. Therefore, we controlled for effects of RIP-Cre presence on glucose homeostasis across littermate mice to ensure our conclusions are not influenced by non-specific effects. To ensure reproducibility, physiological, cellular, and molecular phenotypes from these experiments were replicated in orthogonal mouse and human genetic models. ChIP-sequencing and single-cell sequencing / proteomic experiments were not designed to capture temporal (time-of-day) dynamics. Ligand–receptor interactions were predicted from probabilistic transcriptomic analyses, not experimentally determined. To ensure our conclusions from pharmacologic agents are free from off-target effects, we validated them by genetic manipulation of their targets.

## Methods

### Developmental stage, tissue, and disease state specificity analysis

We analyzed RNA-sequencing data from β cells in humans^40^, pigs^39^, rats^38^, and mice^37^ to compare *Dec1*’s developmental stage specificity to that of adult-restricted transcription factors known to regulate human β-cell functional maturation. Specificity of the expression of a given gene to the fetal or adult stage was quantified as the fraction of the gene’s cumulative expression across fetal and adult stages represented in that stage (i.e. its fractional expression level). To assess *Dec1*’s tissue expression pattern, we queried its expression across mouse tissues and cell types from the Tabula Muris single-cell RNA-sequencing project^74^. To correlate human DEC1 levels with β-cell function in diabetic donors, we compared paired islet functional data and single-cell RNA-seq from type 1 and 2 diabetic, islet autoantibody-positive, and non-diabetic donors to the Human Pancreas Analysis Program obtained from the PANC-DB database^75,76^. All public datasets accessed are listed in Table S1.

### Animals

All animal care and use procedures followed the guidelines of the Institutional Animal Care and Use Committee of the University of Pennsylvania in accordance with the guidelines of the NIH. Mice were generated on the C57BL/6 background, maintained as heterozygotes, and housed in a 12-hour light:12-hour dark cycle (lights on at 6am [zeitgeber time 0, ZT0] and lights off at 6pm [ZT12]) with continuous access to regular chow and water, unless otherwise noted. Male *Dec1^-/-^*and *Dec1^+/+^* littermates used in this study were generated by intercrossing *Dec^+/-^* heterozygotes (RRID:IMSR_JAX:029732). *Dec1^f/f^*mice^43^ (kindly provided by Dr. Lino Tessarollo, National Institutes of Health) were crossed to *InsCre* mice^73^ (kindly provided by Dr. Douglas Melton, Harvard University) to produce *InsCre Dec1^f/+^* heterozygotes, which were then intercrossed to generate the male *InsCre Dec1^f/f^* mice and littermate controls used in this study. Female mice were excluded due to sexual dimorphism of glucose tolerance to distinguish genotype-and time-of-day-dependent effects from confounding hormonal oscillations.

### Primary human samples

Primary human adult islets were obtained from the Integrated Islet Distribution Network (IIDP), the Alberta Diabetes Institute Islet Core (ADI), the UCSF Human Islet Production Core, and the Human Pancreas Analysis Program (HPAP). The cadaveric donors had no history of diabetes and spanned a range of ages and health parameters:

Donor 1: HPAP-200 (HPAP), 51-year-old male, with a BMI of 26.6 and HbA1c 6.0% Donor 2: R591 (ADI), 56-year-old male, with a BMI of 23.1 and HbA1c 5.6%

Donor 3: RHIP-157 (UCSF), 56-year-old female, with a BMI of 24.9 and HbA1c 5.5% Donor 4: SAMN50439434 (IIDP), 21-year-old male, with a BMI of 26.9 and HbA1c 5.1% Donor 5: SAMN39643547 (IIDP), 49-year-old male, with a BMI of 20.9 and HbA1c 5.2% Donor 6: SAMN49690539 (IIDP), 44-year-old female, with a BMI of 33.6 and HbA1c 5.7% Donor 7: RHIP-166 (UCSF), 46-year-old female, with a BMI of 35.8 and HbA1c 5.9% Donor 8: ICRH174 (UCSF), 35-year-old male, with a BMI of 26.5 and HbA1c 5.8%

Donor anonymity was preserved, and donor 1 was used to profile DEC1 protein expression; donors 2-7 for sgRNA knockout experiments; and donor 8 for siRNA knockdown experiments.

All methods involving human cells were approved by the University of Pennsylvania IRB and ESCRO committees.

### Immunostaining

Immunostaining was performed as described previously^36^. Mouse pancreata were fixed in 4% paraformaldehyde overnight at 4°C, dehydrated using increasing concentrations of ethanol, and maintained in 100% ethanol until embedding. Fixed tissue was sectioned by the Penn Molecular Pathology and Imaging Core (RRID: SCR_022420). Sections were deparaffinized using Histoclear (Thermo Fisher Scientific; C78-2-G), submerged in 1X Citrate buffer (Sigma Aldrich; C9999-1000ML) for antigen retrieval, and placed in a pressure cooker for 20 minutes at 90°C. Slides were blocked with 5% Donkey Serum (Jackson Immunoresearch; 017-000-121) and PBST (PBS and 0.3% Triton X-100 [Sigma Aldrich]) for 1 hour, then incubated with PBST-diluted primary antibodies overnight at 4°C: insulin (Agilent; IR00261-2), glucagon (Santa Cruz; SC-514592), Ki67 (Abcam; ab16667), DEC1 (Novus Biologicals; NB100-1800), and/or MAFA (Abcam; ab17976). On the following day, slides were washed twice with PBST and incubated with secondary antibodies at 1:500 dilution in PBST for 1 hour protected from light. For imaging, slides were mounted with Prolong gold Antifade mount and with DAPI (Thermo Fisher Scientific; P36935) and covered with coverslips. Isolated mouse islets were fixed in 4% paraformaldehyde, embedded in agarose gel, and maintained in 100% ethanol, then sectioned by the Penn CVI Histology Cooperative as previously described^77^. Sections were stained using the following antibodies: insulin (Agilent; IR00261-2), PINK1 (Proteintech; 23274-1-AP), PARKIN (Proteintech; 66674-1-Ig). All Images were taken using a Carl Zeiss microscope (LSM 510) at 20x magnification. To analyze PINK1-PARKIN colocalization, the Mander’s coefficient was determined using the JACoP plugin in FIJI.

### Pancreas morphometry

Pancreas morphometry was performed as described previously^78^. ImageJ (NIH) was used to evaluate islet area, α and β cell relative proportions, and β cell proliferation and survival. Islet area was calculated per mm^2^ of pancreatic section area, defined by the total DAPI-positive area. α and β cell proportions were quantified as the fraction of glucagon+ and insulin+ cells per islet, respectively. To quantify proliferating β cells, we calculated the ratio of insulin+ Ki67+ cells to total insulin+ cells within islets. To assess β-cell survival, we calculated the ratio of insulin+ TUNEL+ to total insulin+ cells within islets using a Biotium kit (CF ® 488A TUNEL Assay Apoptosis Detection Kit; 30063) following the manufacturer’s directions.

### Blood glucose and plasma insulin / C-peptide measurements

Blood glucose levels were measured via tail vein bleed using a OneTouch Ultra2 glucometer (LifeScan). Plasma samples were collected via tail vein bleed into a Microvette CB300 (Sarstedt, Inc.) capillary tube. Insulin and C-peptide concentration were quantified using a Mouse Ultrasensitive Insulin ELISA kit (ALPCO Diagnostics; 80-INSMSU) and a Mouse C-peptide ELISA kit (ALPCO Diagnostics; 80-CPTMS-E01) within 24 hours of sample collection.

### Glucose and insulin tolerance tests

Glucose tolerance tests were conducted in mice fasted for 14 hours. Blood glucose levels were measured as indicated above following intraperitoneal injection of room-temperature D-(+)-glucose in 0.9% sterile saline at 2g/kg of body weight. Plasma insulin was measured as indicated above. Insulin tolerance tests were performed after a 4-hour fast, followed by intraperitoneal injection of 4°C insulin (*Humulin R*, Eli Lilly) in 0.9% sterile saline at 1U/kg of body weight.

### Indirect calorimetry and body composition

Indirect calorimetry measurements were collected as described^79^ with mice housed at 23°C in Promethion indirect calorimetry cages (Sable Systems) with free access to food and water. Mice were first assayed under regular light-dark cycles, with a 6am-6pm light photoperiod. The first full 24-hour day was considered an acclimation period, and the following 3 days were used for data collection. Lights were then turned off in the chambers for dark-dark cycle measurements over 7 days, with food shaken down in days 2 and 4 at ZT4, to detect self-sustained behavioral rhythms. Body mass was measured before and after the experiment, and the average was used for regression analyses. Body mass composition was assessed by magnetic resonance imaging with an EchoMRI instrument before placing mice into metabolic cages. CalR analysis (RRID:SCR_015849) was performed as described^79^. For measurements not associated with mass, the difference between groups was analyzed by a one-way ANOVA. Mass-dependent variables were analyzed by analysis of covariance, including an interaction term when genotype × body mass interactions were significant.

### CRISPR-Cas9-mediated *DEC1* knockout in human islets

CRISPR-Cas9 RNP complexes were assembled by incubating 1.25ug of Cas9GFPPRO (Sigma-Aldrich) with 3 sgRNAs (0.65ug each) at room temperature for 15min in 5ul Gene Editing buffer (NeonTM, Thermo Fisher Scientific). *DEC1*-targeting sgRNAs were designed using the E-CRISPR^80^ and CRISPick (RRID: SCR_025148) CRISPR design tools. Chemically modified sgRNAs (with 20-O-Methyl at 3 first and last bases, and 30 phosphorothiate bonds between first 3 and last 2 bases) were purchased from Synthego. CRISPR/Cas9 RNP complexes were assembled by incubating 1.25ug of Cas9GFPPRO with three sgRNAs (0.65ug each) at room temperature for 15 min in 5 ul Gene Editing buffer (NeonTM, Thermo Fisher Scientific). Human islets were dispersed into single cells by enzymatic digestion (Accumax, Invitrogen). We first resuspended 150,000 cells in 5ul Gene Editing buffer/ electroporation pulse. CRISPR-Cas9 RNP complexes were combined with the cells and electroporated in 10ul tips of the NeonTM transfection system (Thermo Fisher Scientific). The electroporation conditions consisted of three 10ms pulses of 1600V. Immediately after pulses, cells were transferred into ultra-low attachment 24 well plates (Corning; 3473) in islet culture medium consisting of RPMI 1640 (Gibco), 2.25 g/dL glucose, 1% penicillin/streptomycin (v/v, Gibco), and 10% fetal bovine serum (HyClone). Electroporated islet cells were cultured in ultra-low attachment plates for 5 days to enable reaggregation into pseudoislets before downstream assays. Genomic DNA was extracted using a Quick-DNA kit (Zymo Research; D3021) as per the manufacturer’s protocol and quantified using NanoDrop. NanoDrop.

### Beta-TC-6 and MIN6 cell culture

All cells were maintained in a 37°C 5% CO_2_ incubator, grown as adherent monolayers, and passaged at ∼70–80% confluence. Mouse insulinoma Beta-TC-6 cells (ATCC; CRL-3605) were cultured in DMEM supplemented with 15% FBS, 1% L-glutamine, and 1% penicillin-streptomycin. All experiments were performed at passage numbers below 20. Mouse insulinoma MIN6 cells (AddexBio; C0018008) were cultured in DMEM (ATCC; 30-2002) supplemented with 15% FBS, 1% Antibiotic-Antimycotic, and 0.05 mM 2-Mercaptoethanol. All experiments were performed at passage numbers below 40. To generate pseudoislets, MIN6 single-cell suspensions were seeded at a density of 6 x 10^5^ cells per mL into wells of ultra-low attachment 6-well plates (Corning; 3471). Plates were placed on an orbital shaker and continuously agitated at 300 rpm. The resulting MIN6 pseudoislets were synchronized by a media shock using complete media 3 days post-seeding, followed by harvesting for downstream assays 4 days post-seeding.

### ChIP-sequencing

Chromatin immunoprecipitation-sequencing was conducted as described^36^, by incubating Beta-TC-6 cells with DEC1 (Novus Biologicals; NB100-1800) antibody and Beta-TC-6 cells expressing a DEC2-V5 plasmid (GeneCopoeia, Inc. # EX-A4343-LX304) with anti-V5 (abcam; ab15828) antibody. Sequencing reads were mapped using STAR^81^ with default parameters.

Regions enriched for uniquely-mapped reads relative to input sample were called as binding peaks using MACS^82^ with default parameters and “--pvalue=1e-9 --keep-dup=1”, followed by filtering against overlap with problematic regions of the genome^83^. Peaks from BMAL1, CLOCK, and PDX1 ChIP-sequencing in Beta-TC-6 cells^29^ were called in the same manner and concatenated and merged with DEC1 and DEC2 peaks using BEDTools^84^ to obtain a unified peak catalog. Reads in peaks from the unified catalog were quantified using HTSeq^85^ in ‘‘-m union’’ mode, and normalized as counts per million mapped reads (CPM). K-means clustering of peaks by their read content across the DEC1, DEC2, BMAL1, CLOCK, and PDX1 ChIP-sequencing datasets was conducted in R (kmeans function) as described^36^ to identify binding clusters.

DEC1 and NCOR1^86^ ChIP-seq peaks were intersected by their genomic coordinates, and overlap was defined as ≥1 bp shared between peak regions. Statistical significance was assessed using a permutation-based overlap test (regioneR^87^, 2,000 permutations on the mm9 genome), testing if the observed number of overlapping peaks exceeds random expectation.

### Motif enrichment

DNA sequences of DEC1 and DEC2 binding peaks were searched for enriched motifs from a database of vertebrate transcription factor recognition motifs^88–90^ using MEME-ChIP ^91^ under Discriminative mode, which implements CentriMo^92^ to identify centrally enriched motifs and tests for similarity to known motifs using Tomtom^93^. Markov models of sequence nucleotide composition were used as the background model for motif matching to normalize for biased distribution of individual letters in the examined sequences. We used FIMO^94^ as described^36^ to search regions of interest (e.g., clusters of peaks from the unified peak catalog) for matches to the database of vertebrate transcription factor recognition motifs.

### Pathway enrichment

Gene lists ranked by a feature of interest (e.g., expression change,-log10 p-value of rhythmicity) were analyzed for enrichment of gene sets grouped by functional annotations using Metascape^95^ with default parameters, followed by manual curation of a representative term for each ranked group of enriched annotation terms. Genomic regions of interest (e.g. clusters of peaks from the unified peak catalog) were analyzed for enrichment of associated gene functional annotations using HOMER^96^ with default settings, followed by p-value based ranking and manual curation to prune redundant terms.

### RNA-sequencing and differential expression

Isolated islets were sampled, dissociated, and subjected to RNA-sequencing at 4-hour intervals 26 to 50 hours post-isolation. RNA-sequencing was conducted as described^36^, Sequencing reads were mapped using STAR^81^ with default parameters. Read counts per gene were determined using HTSeq^85^, and normalized using DESeq^97^. We compared gene expression distributions (regardless of rhythmicity) between *Dec1 βKO* and wild-type islets using dryR^51^ (see below) to identify genes with differences in mean expression level, and considered genes with DESeq p <0.05 in both the islet-level 24-hour time series and the β cell-level scRNA-seq data (see below) to be differentially expressed.

### Rhythmicity analysis

RNA-sequencing data from *Dec1 βKO* and wild-type islets was analyzed for 24h rhythmicity using dryR^51^, which employs a Bayesian information criterion for rhythm and mean expression model selection, considering expressed only genes with cumulative expression >10 normalized counts across the 24-hour time series of either genotype. Default parameters were used to detect and estimate changes in amplitude, peak phase, and mean expression level between the two 24-hour series of N =7 time-points with n =4 replicate measurements each. To establish statistical confidence for a single condition, we used the “dryseq_single” function to compute a p value for each gene. To quantify and compare rhythm phase shifts between *Dec1 βKO* and wild-type islets for gene sets driving enriched biological themes, we used PSEA^98^, employing a significance threshold of q <0.05 against a uniform phase distribution. Other features of interest (e.g. mitochondrial structural and functional measurements) were analyzed for 24h rhythmicity using RAIN^99^, which employs non-parametric Mann-Whitney U tests to compare the ranks of measured values against those of alternative waveforms without assumptions about waveform shape or symmetry and computes a Benjamini-Hochberg corrected p value for each feature.

Default parameters and “method = independent” were used to estimate peak phase, shape, and significance from the time series sampling independent populations. Comparable trends and statistical significance estimates were obtained using parametric methods such as harmonic regression. For plotting, time series measurements were detrended by subtracting a linear regression fit, maintaining relative scaling proportions by considering the means and SEMs of both genotypes.

### CRISPR-Cas9-mediated *Dec1* knockout in MIN6 cells

*Dec1-*knockout MIN6 cells were generated using a plasmid-based CRISPR-Cas9 approach. MIN6 cells were transfected with a *Dec1*-targeting CRISPR-Cas9 knockout plasmid (Santa Cruz Biotechnology; sc-423203) plus a donor plasmid for homology directed repair (Santa Cruz Biotechnology; sc-423203-HDR) following the manufacturer’s instructions. 72h after transfection, cells were subjected to puromycin selection at 2ug/mL to eliminate untransfected cells, maintaining selection for 7 days. Genome editing at the *Dec1* locus was confirmed by PCR-mediated amplification of the targeted region followed by Sanger sequencing. DEC1 protein loss was validated by western blot analysis.

### DEC1 co-immunoprecipitation followed by mass spectrometry

Cell lysates from *Dec1^-/-^* and *Dec1^+/+^*MIN6 cells were prepared in RIPA buffer (CST; 9806S) supplemented with protease and phosphatase inhibitors and clarified by centrifugation (14,000 × g, 15 min, 4 °C). Protein concentrations were determined using a BCA assay (Thermo Fisher Scientific; 23225). For immunoprecipitation, Dynabeads Protein G (Thermo Fisher Scientific; 10003D) were incubated with 5 µg anti-DEC1 antibody (Novus Biologicals; NB100-1800) or control IgG (CST; 2729S) for 10 min at room temperature with gentle rotation. Antibody-bound beads were subsequently washed three times with PBS-T and incubated with 300 µg clarified protein lysate overnight at 4 °C with continuous rotation. Beads were then washed with PBS-T (0.1% Tween-20) and bound proteins were eluted in LDS sample buffer at 70 °C for 10 min.

Eluates were resolved briefly on 10% SDS–PAGE gels (∼0.5 cm), stained with Coomassie blue, and gel regions containing proteins were excised for in-gel trypsin digestion. Tryptic peptides were analyzed by LC–MS/MS using an Orbitrap Astral mass spectrometer coupled to a Vanquish Neo UPLC (Thermo Fisher Scientific). Peptides were separated on a C18 nanocapillary column using a 40-min gradient (5–80% acetonitrile with 0.1% formic acid) at 300 nL min⁻¹. Data were acquired in data-independent acquisition (DIA) mode. Raw data were processed with DIA-NN (v2.2.0)^100^ against the UniProt mouse proteome (UP000000589).

Searches allowed two missed tryptic cleavages and common variable modifications. Contaminants and proteins supported by a single peptide were removed. Protein abundances were normalized across conditions and differential enrichment was assessed using LFQ Analyst^101^ with Benjamini–Hochberg correction; proteins with fold change > 2 and adjusted p < 0.05 were considered significant.

### Mouse islet isolation

Islets were isolated at ZT2 from adult mice as previously described^102^, via *in situ* pancreas perfusion with 0.8 mg/ml collagenase P (Roche; 11-213-873-001) followed by pancreas digestion and islet purification using Histopaque 1077 and 1119 (Sigma). Islets were filtered through a 250 μm mesh filter, collected in a 40 μm mesh filter, and manually picked with a 200 µL micropipette, followed by overnight recovery in a 37°C 5% CO2 incubator prior to downstream experiments. Temporal patterns in downstream experiments were analyzed relative to hours post-isolation, reflecting the cell-synchronizing effect of islet isolation.

### *In vitro* glucose-stimulated insulin secretion assays

Insulin secretion assays were conducted in mouse islets after an overnight recovery from isolation, in human primary islets after an overnight recovery from receipt, and in MIN6 pseudoislets as previously described^36^. 20 size-matched islets were allotted into four groups, with one group used to determine insulin content and three groups used for triplicate GSIS measurements. For insulin content determination, islets were directly dissolved in M-PER protein extraction reagent (Thermo Fisher Scientific; 78501). For GSIS measurements, isles were washed with Krebs buffer containing basal (0.5/2.8mM) glucose and loaded into 24-well plate inserts (Millicell Cell Culture Insert; PIXP01250), followed by pre-incubation in Krebs buffer with the same basal (0.5/2.8mM) glucose level for 1h to remove residual insulin. Subsequently, islets were sequentially challenged with Krebs buffer containing indicated glucose concentrations with or without insulinotropic agents, with a 1h incubation for each treatment.

Insulinotropic agent dosages were: 10µM Ionomycin (STEMCELL; 73722); 10µM Forskolin (Stemolecule; 04-0025); 1mM 8-Br-cAMP (VWR; 80050-494); 1mM Carbachol (Santa Cruz; sc-202092); 10µM PMA (Santa Cruz; sc-3576B); 10µM Dorzagliation (Selleckchem; S6921); 10µM TEPP-46 (ML265) (Selleckchem; S7302) or 3µM Cyclosporin A (CSA, Enzo; BML-A195-0100). For chronic pharmacologic GSIS rescue, 1mM dichloroacetate (DCA, Selleckchem; S8615) was used to treat overnight-recovered mouse islets for 24 hours before the GSIS assay. All incubations were conducted in a 37°C 5% CO_2_ incubator, with supernatant samples collected after each incubation. After GSIS assays, islets were dissociated using TrypLE Express (Life Technologies) and cell counts were obtained using an automated cell counter (Beckman Coulter; Vi-Cell).

Dynamic GSIS assays of handpicked, size-matched islets were conducted in triplicate as previously described^36^ using an automated Perifusion System (BioRep). Chambers were first perifused with Krebs buffer containing basal (0.5/2.8mM) glucose for 1h at a flow rate of 100ul/min and sequentially with the indicated solutions for the indicated durations. Dynamic insulin secretion measurements were normalized to the mean of the first (basal) incubation.

### Single-cell RNA and ATAC-sequencing

Islets were isolated at ZT2 and dissociated using Accutase (StemCell Technologies), followed by single-cell capture, lysis, and parallel RNA and ATAC library preparation with the 10x Chromium system (10x Genomics) using Single Cell 3’ Gene expression (v3) and ATAC reagents following the manufacturer’s instructions. Libraries were sequenced on a NovaSeq 4000 instrument (Illumina) and aligned to the mm10 reference genome assembly using Cell Ranger (10x Genomics) RNA/ATAC pipelines. Mapped RNA reads were processed as described previously^103^ according to guidelines from^104^ using Seurat V3^57^. ATAC reads were processed as described^105^, filtering against low-quality fragments and excluding regions within non-standard chromosomes or with extremely high (top 2%) nucleosome signal, to minimize noise and artifacts. Principal component analysis was performed on log-scaled counts for the top 2000 highly variable genes, and uniform manifold approximation and projection (UMAP) was computed with the top 50 principal components. Clusters were inferred using the Seurat “FindClusters” function adjusting for cluster resolution to optimize balance between cluster granularity and separation. Cross-modal integration of RNA and ATAC data was performed with Seurat^57^ using the “FindTransferAnchors” and “TransferData” functions with a Canonical Correlation Analysis (CCA) dimension reduction method to transfer cluster identities from RNA to ATAC data. Cell clusters were then annotated to cell types based on established islet cell identity markers^57^.

Gene expression / promoter accessibility were computed for each cell type as expected counts within gene models / promoter-proximal (TSS ± 2kb) regions using RSEM^106^. Differential expression / promoter accessibility was evaluated in β-cells as described^107^ using edgeR^108^.

Uniquely-mapped reads were removed for duplicates and used for peak calling as described ^36^ using MACS2^82^ with default parameters and ‘‘–broad –qvalue 0.01’’, followed by filtering against very small (<20bp) or large (>10kb) peaks and against overlap with problematic regions of the genome^83^. Differential peak counts were computed using DiffBind^109^ with default settings.

### Single-cell proteomics

Islets were isolated at ZT2 and single-cell samples were prepared using the droplet nPOP method as detailed in^56^. Sets of 14 single cells were run with a carrier sample of 100 cells of bulk digest labeled with TMT 126C and a 5-cell reference labeled with TMT 127N prepared via mPOP sample preparation^110,111^. Channels 127C and 128N were not used due to isotopic contaminants from the carrier channel.

For LC/MS analysis, samples were separated via online nLC on a Dionex UltiMate 3000 UHPLC; 1 μl out of 1.1 μl of sample was loaded onto a 25 cm × 75 1 μm IonOpticks Aurora Series UHPLC column (AUR2-25075C18A). Buffer A was 0.1% formic acid in water and buffer B was 0.1% formic acid in 80% acetonitrile / 20% water. Samples were analyzed by a Q-Exactive mass spectrometer (Thermo Scientific). After a precursor scan from 450 to 1600 m/z at 70,000 resolving power, the top 7 most intense precursor ions were isolated for MS2 analysis via a 0.7 Th isolation window with a 0.3 Th offset. These ions were accumulated for at most 500 ms before being fragmented via HCD at a normalized collision energy of 33 eV (normalized to m/z 500, z = 1). The fragments were analyzed at 140,000 resolving power.

Single cell MS data was preprocessed via the SCoPE2 data pipeline^112,113^. Cell types were then assigned via label transfer from scRNA data performed with LIGER^114^. Statistical tests between genotypes were performed via a two-tailed t-test comparing protein expression between groups of single cells, using the Benjamini-Hochberg adjustment to correct for multiple hypotheses testing. Data and code are reported in accordance with community guidelines^115^ and are available on github (https://github.com/SlavovLab/SC_Pancreas_KO).

### Ligand-receptor interaction analysis

Single-cell RNA-sequencing data was analyzed using CellChat^116^ (v.2.1.2) to infer intercellular communication changes in *Dec1-βKO* vs. wild-type islets. CellChat infers probabilistic cell–cell communication networks from scRNA-seq data through statistical filtering (overexpression tests), database-encoded biology (complexes + cofactors), mass-action modeling, and network-based smoothing/regularization. Enriched signaling genes and ligand-receptor interactions in each genotype were identified using the “identifyOverExpressedGenes” and “identifyOverExpressedInteractions” functions with default settings. A protein-protein interaction network was applied to smooth predictions. Communication probabilities were computed with “computeCommunProb” and filtered to exclude interactions involving <10 cells. Pathway-level predictions were refined using “computeCommunProbPathway”, and communication networks were aggregated with “aggregateNet”. Network centrality was assessed via “netAnalysis_computeCentrality”, highlighting key signaling roles such as influencers or receivers within each genotype. To identify the top 10 differentially active pathways (p <0.01 for change in communication probability between *Dec1-βKO* and wild-type), the overall information flow through signaling pathways across all cell types was ranked using the “rankNet” function.

We then selected ligand-receptor pairs where the ligand is significantly altered (p <0.05) in β1 cells (β cells with higher expression of insulin and maturity markers) and calculated the log2 fold change in communication strength between *Dec1-βKO* vs. wild-type cells to quantify the change in communication flow.

### Cellular and mitochondrial energetics

Imaging of stained mitochondria was conducted in mouse islets after an overnight recovery from isolation. All incubations were performed at 37°C with 5% CO_2_. 30 islets were hand-picked, dissociated using TrypLE Express (Life Technologies), and the single cells seeded onto glass-bottom dishes (Cellvis; D29-14-1.5-N) pre-coated with 500ng/ml concanavalin A (Sigma; C2010) and incubated for 30 minutes to facilitate attachment. Plated cells were incubated with freshly prepared 500nM PKmito ORANGE FX (Cytoskeleton; CY-SC054) in RPMI-1640 medium (Life Technologies; 11875-093) for 2 hours, then washed and fixed following the manufacturer’s instructions, followed by immunostaining with PINK1 (Proteintech; 23274-1-AP) and(Proteintech; 66674-1-Ig) as described above Z-stack images were acquired with a 63x magnification lens using a Leica confocal microscope (Stellaris 5) and pre-processed by deconvolution using the PSF generator and DeconvolutionLab2 plugins in FIJI^117^. The Mitochondria Analyzer tool^118^ was then used to skeletonize the deconvoluted z-stack images and quantify morphometric parameters.

Flow cytometry analysis of mitochondrial content and activity, cellular reactive oxygen species, and glucose uptake was conducted in mouse islets dissociated as above. Single cell preparations were stained with freshly prepared solutions of either MitoTracker Green FM (Invitrogen; M46750; 1:2000) and 5µM CellROX Deep Red (Invitrogen; C10422), 50nM TMRE (Abcam; ab113852), or 20uM 2-NBDG (Invitrogen; N13195) in RMPI-1640 medium and incubated for 30 minutes at 37°C with 5% CO_2_. Samples were then washed twice with 1% BSA in 1x PBS and filtered through a 40µm nylon mesh into flow cytometry tubes (STELLAR SCIENTIFIC; FSC-9005). Flow cytometry was conducted with a BD Accuri C6 Plus flow cytometer and analyzed using FlowJo v10.8.1 (BD Life Sciences).

### Metabolic flux analysis

The Agilent Seahorse XFe96 Spheroid FluxPak (Agilent; 102905-100) and Seahorse XFe96 Analyzer were used to measure the oxygen consumption rate (OCR) and extracellular acidification rate (ECAR) of mouse islets after an overnight recovery from isolation. Subsequent incubations were performed in a 37°C CO_2_-free incubator. The day before the experiment, sensor cartridge wells were rehydrated by incubating with 200µL of Agilent calibrant overnight, and the experimental wells of the spheroid plate were pre-coated with 30µL of 100 ng/mL poly-D-lysine (Sigma; P7280). Injection basal media (Agilent; 103335-100), wash media (basal media with 0.5% FBS), and assay media (basal media with 0.5% FBS and 1mM glucose) were freshly prepared and stored at 4°C protected from light. On the day of the experiment, all media were adjusted to pH 7.4 and warmed and kept at 37°C. Islets were hand-picked, incubated in wash media for two sequential 15-minute washes at room temperature, and transferred to each well of the spheroid plate pre-loaded with 175µL of assay medium (12 islets in a 30µL volume per well) using an islet-loading mold by gravity. The spheroid plate with loaded islets was then equilibrated for 1hr at 37°C. The Seahorse XFe96 Analyzer was calibrated with the cartridge plate for 15 minutes before data acquisition. For mitochondrial stress assays, 20µL compounds were sequentially injected to achieve final concentrations of 16.7 mM glucose supplemented with 0.5 mM glutamine and 3.5 mM amino acid mix^119^ (Table S6), 5 µM oligomycin (Sigma; O4876), 2 µM FCCP (Sigma; C2920), and 5 µM rotenone (Sigma; 557368) plus 5 µM antimycin A (Sigma; A8674). Assay cycles were performed as follows: 6 cycles for basal OCR/ECAR measurements; 4 for glucose plus amino acids; 5 for oligomycin; 4 for FCCP; and 6 for rotenone plus antimycin A. For glycolytic flux assays, 20µL compounds were sequentially injected to achieve final concentrations of 3.5 mM amino acid mix, 16.7 mM glucose, 50 mM 2-deoxy-glucose (Selleckchem; S4701), and 5 µM oligomycin. Assay cycles included 6 cycles for basal OCR/ECAR measurement; 4 for amino acids; 5 for glucose; 4 for 2-DG; and 6 for oligomycin.

Each cycle consisted of 2 minutes of mixing followed by 3 minutes of measurement. Data were normalized to islet equivalents (IEQ) using images captured at the end of the experiment with a Cytation 5 instrument (Agilent).

### Calcium influx

Extracellular calcium influx dynamics were assayed in mouse islets after an overnight recovery from isolation as previously described^120^. Islets were washed two times with Krebs buffer, followed by staining with 16.7μM Fura2-AM (Life Technologies; F-F1225) in Krebs buffer supplemented with 5mM glucose for 45 minutes at 37°C. Islets were then transferred to a perifusion chamber on a homeothermic platform, perifused with 37°C Krebs buffer at a flow rate of 1mL/min. and imaged with an inverted Zeiss microscope (Axio Observer.Z1). After imaging in the absence of substrate, a physiological amino acids mixture consisting of 0.5 mM glutamine and 3.5mM amino acid mix (Table S6) alone and with increasing glucose levels (3mM, 16.7mM) were sequentially applied, followed by washing all substrates away and applying KCl (30mM). Intracellular Ca^2+^ was measured by dual-wavelength fluorescence microscopy using the Zeiss AxioVision system as previously described^121^.

### Western blotting

Total proteins were extracted from human primary islets or MIN6 pseudoislets by RIPA lysis buffer (Cell signaling; 9806S). Protein concentration was measured by BCA Protein Assay Kit (Thermo Fisher Scientific; 23225). 30μg of proteins were separated on 4-15% precast protein gels (Bio-Rad; 4568083) and transferred to nitrocellulose membranes (Bio-Rad; 1620115) or PVDF membranes (Bio-Rad; 1620177). The membranes were blocked in 5% milk or 5% BSA for 1 hour and incubated overnight at 4°C with the following antibodies overnight at 4 °C: Vinculin rabbit monoclonal antibody (Cell signaling; 13901T) diluted to 1:1000 as the loading control; phosphoSer616-DRP1 antibody (Cell signaling; 3455S) diluted to 1:1000; total OXPHOS antibody cocktail (Abcam; 110413) diluted to 1:1000; NCoR1 antibody (Cell signaling; 5948S) diluted to 1:1000; and DEC1 antibody (Novus; NB100-1800) diluted to 1:500.

After washing, the membranes were incubated for an hour at room temperature with anti-mouse HRP-conjugated secondary antibodies (Cell signaling; 7076S) or anti-rabbit HRP-conjugated secondary antibodies (Cell signaling; 7074P2), both diluted to 1:5000. The blots were visualized by Pico chemiluminescent substrate (Thermo Fisher Scientific; 34580) and exposed in an Amersham Imager 600.

### Quantitative RT-PCR

For mouse islets, MIN6 pseudoislets, and siRNA-treated human islets, RNA was isolated and purified using a Direct-zol RNA miniprep kit (Zymo research; R2050). cDNA was synthesized using the HiScript IV all-in-one ultra RT supermix for qPCR (Vazyme; R433-01). Gene expression was assessed via quantitative PCR using PowerUp SYBR green master mix for qPCR (Thermo Scientific; A25742) in a StepOnePlus Real-Time PCR system (Applied Biosystems).

For sgRNA-treated human islets, RNA was isolated and purified using a PicoPure RNA isolation kit (Life Technologies; KIT0204). cDNA was synthesized using a Maxima first strand cDNA synthesis kit (Thermo Scientific; K1641). Gene expression was assessed using TaqMan gene expression mix (Applied Biosystems) in a QuantStudio 3 Real-Time PCR system (Applied Biosystems). All qPCR primers are listed below in the Oligonucleotides table.

### siRNA-mediated knockdown studies

Primary human islets were transfected with a non-targeting control siRNA (Sigma; SIC007) or with pooled siRNA targeting *DEC1* (Sigma; SASI_Hs01_00157761, SASI_Hs01_00157762, SASI_Hs01_00157763). Mouse islets were transfected with the non-targeting control siRNA described above or with pooled siRNA targeting *Pdk4* (Sigma; SASI_Mm01_00053022, SASI, SASI_Mm01_00053023, SASI_Mm01_00053024). Briefly, siRNA was mixed in Opti-MEM medium (Thermo Fisher Scientific, 31985070). In parallel, Lipofectamine RNAiMAX (Thermo Fisher Scientific; 13778075) was diluted in Opti-MEM. The diluted siRNA and Lipofectamine RNAiMAX solutions were combined and incubated at room temperature for 5 min to allow for complex formation. The transfection mixture was then added to islets to reach 150 nM final concentration. Mouse islets were harvested 72 h post-transfection for downstream end-point assays. Human primary islets were synchronized with 10uM Forskolin (Stemgent; 04-0025) 48 h post-transfection, followed by time-series assays performed 16 h after recovery from forskolin shock to measure self-sustained rhythms.

### Overexpression studies

Lentiviral PKM2 overexpression plasmid (GeneCopoeia, Ex-Mm04490-Lv205) or eGFP control plasmid (GeneCopoeia, LT001) were transfected into mitotic Lenti-X 293T cells (Takara Bio, 632180) maintained in DMEM (Life Technologies) in a 37°C 5% CO_2_ incubator. Viral supernatants were collected 48 h and 72 h post-transfection, filtered (Millipore, SE1M003M00), and concentrated by overnight precipitation at 4°C using PEG-it virus precipitation solution (System Biosciences, MSPP-LV810A1). Precipitated viruses were pelleted by centrifugation at 3,000 x g for 30 min at 4°C, resuspended, and stored at-80°C until use for infection. Isolated mouse islets after overnight recovery from isolation were infected with viral particles for 48 h followed by harvesting for downstream assays.

### Transmission electron microscopy

For ultrastructural evaluation by electron microscopy, MIN6 pseudoislets synchronized by a complete media shock were collected at specific post-shock times followed by overnight 4℃ fixation with 2.5% glutaraldehyde, 2.0% paraformaldehyde in 0.1M sodium cacodylate buffer, pH7.4. After subsequent buffer washes, the samples were post-fixed in 2.0% osmium tetroxide with 1.5% K_3_Fe(CN)_6_ for 1 hour at room temperature and rinsed in dH2O prior to *en bloc* staining with 2% uranyl acetate. After dehydration through a graded ethanol series, the samples were infiltrated and embedded in EMbed-812 (Electron Microscopy Sciences, Fort Washington, PA). Thin sections were stained with uranyl acetate and SATO lead and examined with a Talos electron microscope (L120C) fitted with a Ceta 16M digital camera at 5300x and 17500x magnification. ImageJ (NIH) was used to quantify mitochondrial width:height aspect ratios.

### Oligonucleotides

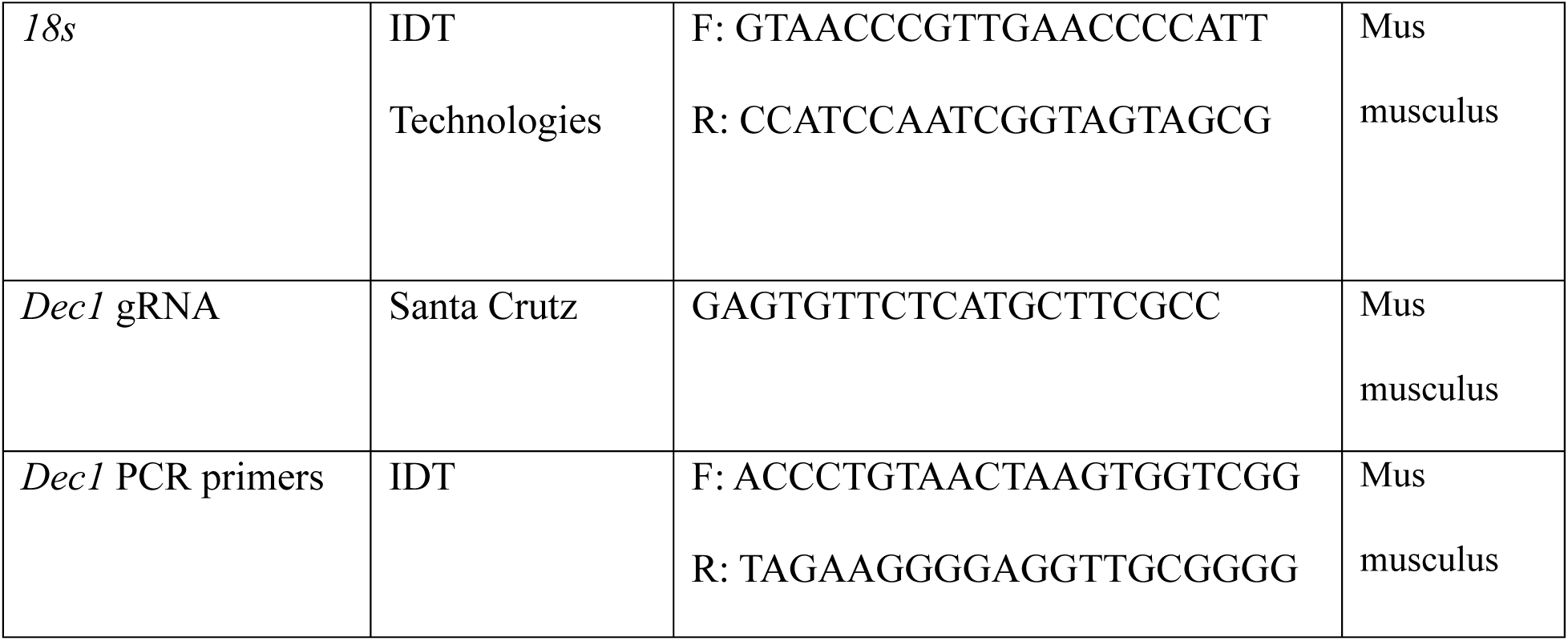
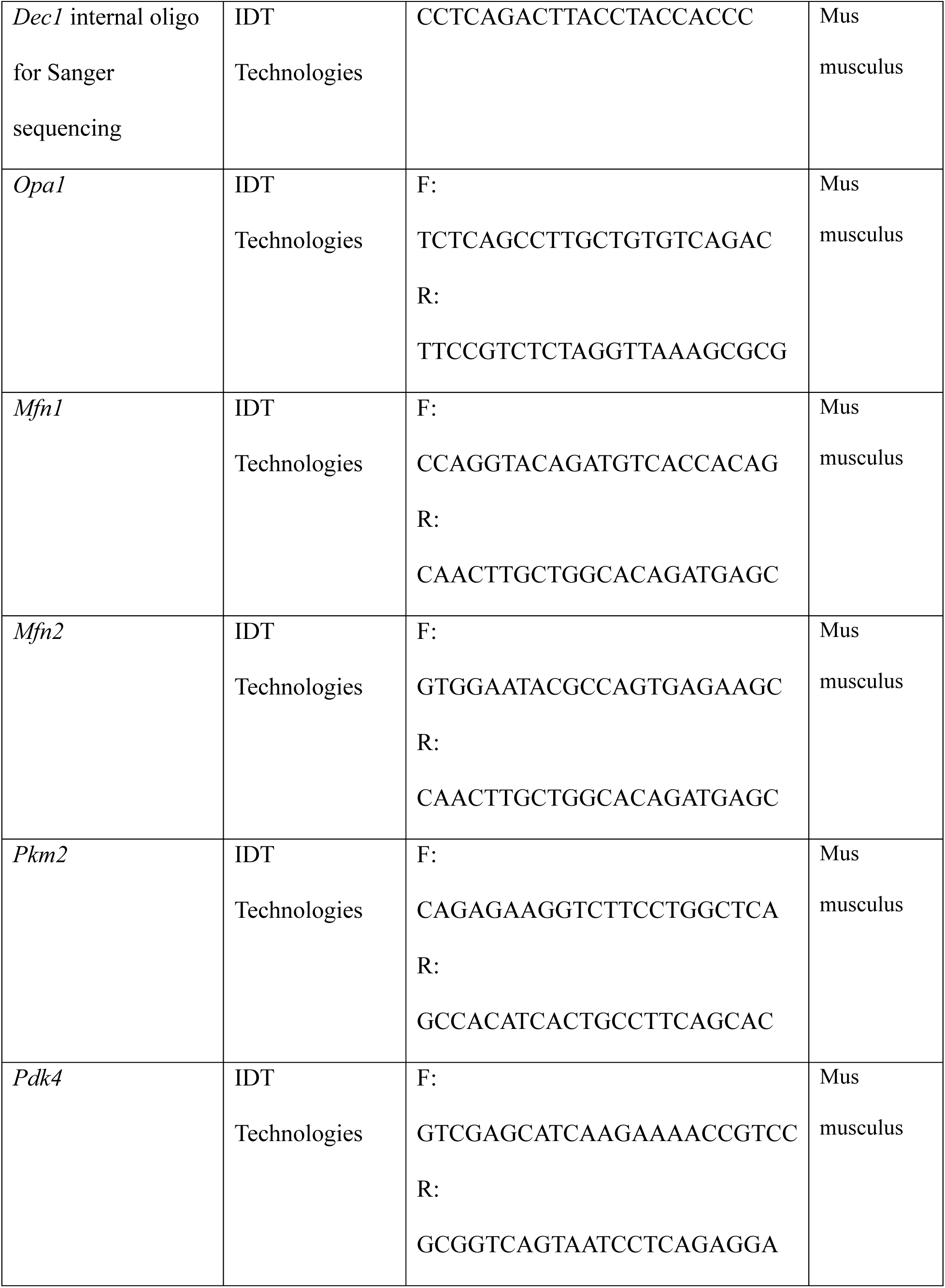
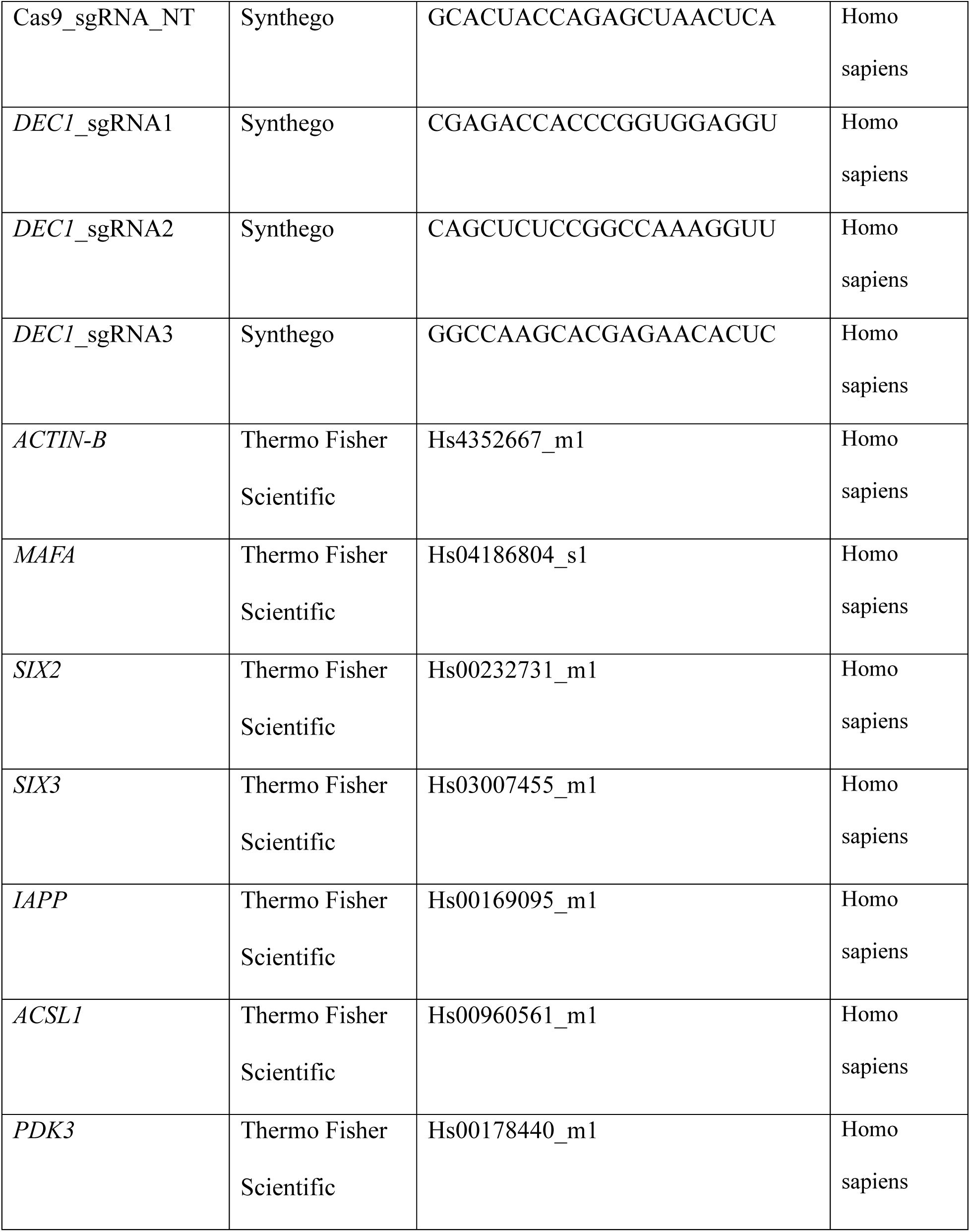

### Rigor and reproducibility

Mice from at least two independent littermate cohorts for each genotype were assayed for each experiment. All mice were housed individually for metabolic assays, with at least 1 week to recover between assays, including light/dark-phase measurements for the same cohort. Mice of all genotypes were assayed in parallel, blinded for genotype whenever possible, to minimize ascertainment bias. Differential gene regulation was validated across genomic/proteomic assays performed on independent mouse cohorts.

### Quantification and statistical analysis

No statistical methods were used to predetermine sample size or remove outliers. To assess the statistical difference between two sets of unpaired data, a Shapiro-Wilk normality test was first performed via the shapiro.test R implementation with default parameters. Normally distributed data was then tested for equality of variance using the Bartlett.test R implementation with default parameters. A two-sided t-test (t.test R implementation with default parameters and var.equal =FALSE or TRUE) was then used to assess confidence on the measured difference of their mean values. For unpaired data that did not follow a normal distribution, we used a non-parametric Wilcoxon rank sum test to determine if they belong to the same parent distribution. For comparisons involving more than two groups, one-way ANOVA was performed for normally distributed data. For experiments involving two independent variables (e.g., genotype and time or treatment), two-way ANOVA was performed. When ANOVA assumptions were not met, a non-parametric Wilcoxon rank sum test was used.

## Acknowledgements

We thank Douglas A. Melton for his generous support during the initial stage of this project; Lan Cheng, Ming Yu, Jasmine Zhao, Biao Zuo, and Anneliese Faustino for reagents and assistance with experiments; Daria Zhadanova, Israeli M. Galicia-Silva, Samuel D. Pollock and Azariah Wise for assistance with data analysis and graphics; the Beth Israel Deaconess Medical Center Metabolic Core, Harvard Medical School Biopolymers Facility, Harvard University Bauer Core sequencing facility, the University of Pennsylvania Laboratory Animal resources, Diabetes Research Center Islet Cell Biology Core, Cell and Developmental Biology Microscopy Core, and Electron Microscopy Resource Lab, and the Wistar Institute Proteomics and Metabolomics Facility for technical support and critical discussions; and Mitchell A. Lazar, Amita Sehgal, Kenneth S. Zaret, Stewart A. Anderson, Hsin Yao Tang and members of the Alvarez-Dominguez laboratory for critical feedback on this manuscript. This work was supported by grants from the NIH (K01DK129442, R35GM157320), the Human Islet Research Network (U24DK104162), and by a pilot award from the Diabetes Research Center at the University of Pennsylvania (P30DK19525) to J.R.A-D., who was also supported by a Howard Hughes Medical Institute Life Sciences Research Foundation Fellowship; and by grants to Douglas A. Melton from the JDRF (5-COE-2020-967-M-N), and the JPB Foundation (award no. 1094).

## Author contributions

Z.G., A.P.M.L., Z.L.G., A.L., Y.X., S.P., J.D., X.L., V.H., Z.C., C.K., M.H., A.V.R., X.W., S.L., L.L., W.Q., and J.RA.-D. performed experiments. M.L., Z.G., A.L., S.J.L., A.B., J.H.A., J.R.S., M.M.C., L.L.M. and J.RA.-D. conducted bioinformatics analyses. Z.G., M.L., A.P.M.L., Z.L.G., C.L., A.N., R.J., N.D., D.A.S., R.B., P.K., N.S., and J.RA.-D. designed the research, interpreted the results, and wrote the manuscript.

## Data and materials availability

Raw and processed sequencing data generated in this study are available at NCBI GEO accession number GSE289105. Proteomic raw data are deposited at MassIVE (MSV000092610).

Processed proteomic data can be accessed at https://github.com/SlavovLab/SC_Pancreas_KO.

## Lead contact and materials availability

Further information and requests for resources and reagents should be directed to the Lead Contact. This study did not generate new unique reagents.

## Supplementary Information

Extended Data Figs 1-8

Supplementary Tables 1-6

Table S1. Datasets used in this study.

Table S2. Unified catalog of DEC1, DEC2, BMAL1, CLOCK, and PDX1 ChIP-sequencing peaks in β-TC-6 cells.

Table S3. Genes that are bound by DEC1 in β-TC-6 cells, and genes that are differentially rhythmic or differentially expressed in *Dec1-βKO* islets.

Table S4. Proteins that are differentially expressed in *Dec1-βKO* islets.

Table S5. Pathway enrichment analysis of DEC1-bound genes that are differentially rhythmic and differentially expressed in *Dec1-βKO* islets.

Table S6. Amino acid mix composition used in this study.

---

**Extended Data Fig. 1.**
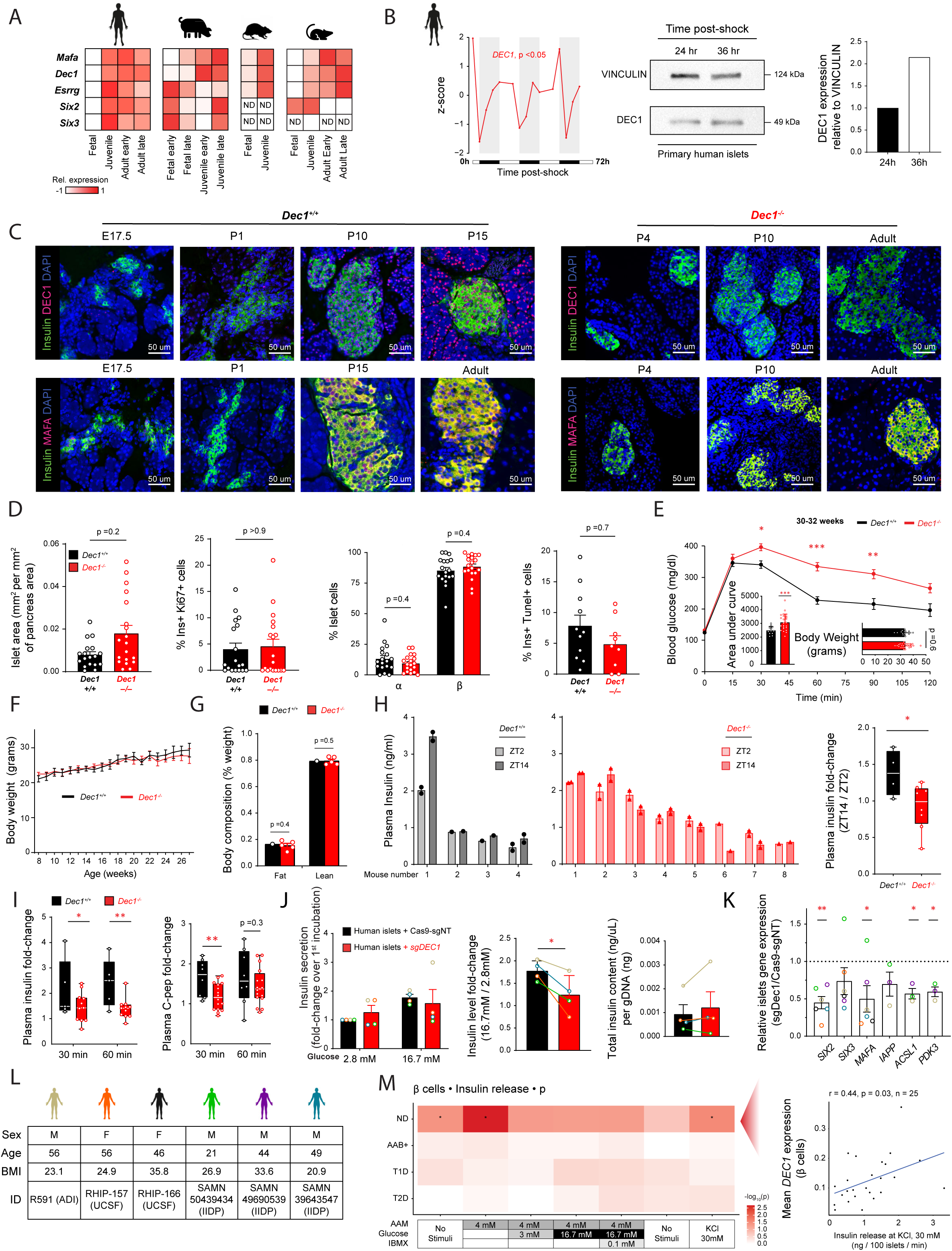
DEC1 defines mature β cells and regulates blood glucose tolerance but not islet formation or body metrics. (**A**) Developmental stage specificity of *Dec1* for adult β cells, compared to adult-restricted transcription factors known to regulate human β-cell functional maturation. Data are average expression from RNA-sequencing of unsynchronized β cells from: human fetal (12-18 gestational weeks, N =6), juvenile (4-19 years, N =3), and adult (24-30 years, N =2; 53-60 years, N =2) donors^40^; pig fetal (embryonic day 40-70, N =11; embryonic day 90, N =2) and juvenile (postnatal day 8, N =2; postnatal day 22, N =3) ages^39^; rat fetal (embryonic day 19, N =3) and juvenile (postnatal week 2, N =3) ages^38^; and mouse fetal (embryonic day 12.5-15.5, N =4), juvenile (postnatal day 16, N =3) and adult (2–3 months and 182 days; N = 4) ages^37^. (**B**) Rhythmic *DEC1* expression in forskolin shock-synchronized primary human islets across a 0-72h time course (left)^36^. Normalized expression from bulk RNA-sequencing is shown as z-scores. Rhythmicity p-value, RAIN. DEC1 immunoblot (middle) at 24h and 36 post-shock and quantification (right). (**C**) Immunofluorescence staining of DEC1 (magenta) or MAFA (magenta), insulin (green) and DAPI (blue) in pancreata from *Dec1^+/+^* and *Dec1^-/-^* littermate mice across developmental stages, from embryonic stage 17.5 (E17.5) to adulthood (2 months), acquired at 20x magnification. (**D**) Quantification of islet area, fraction of proliferating (Ki67+) β cells, cell composition (% α and β cells), and fraction of apoptotic (TUNEL+) β cells across pancreas sections from N=2 *Dec1^+/+^*and 2 *Dec1^-/-^* 2-month-old littermate mice. Data are mean ±SEM. (**E**) Glucose tolerance in 14-hour fasted N =12 *Dec1^+/+^*and 25 *Dec1^-/-^* 8-month-old littermate mice following intraperitoneal glucose administration at ZT2. Data are mean ±SEM. (**F**) Weight of *ad libitum* fed N =16 *Dec1^+/+^* and 12 *Dec1^-/-^* littermate mice at ZT12 monitored from age 8-27 weeks. Data are mean ±SEM. (**G**) Body composition of N =1-5 *Dec1^+/+^*and *Dec1^-/-^* 9-10-month-old littermate mice, determined by magnetic resonance imaging with an EchoMRI instrument. Data are mean ±SEM. (**H**) Plasma insulin in *ad libitum* fed N =4 *Dec1^+/+^* and 8 *Dec1^-/-^*4-5-month-old littermate mice at ZT2 and ZT14. Individual mouse data (left, *Dec1^+/+^*; middle, *Dec1^-/-^*) and fold change (ZT14/ZT2) in plasma insulin (right). Data are mean ±SEM. (**I**) Plasma insulin (left) and C-peptide (right) fold change 30 min and 60 min following intraperitoneal glucose injection at ZT6 in N = 5-8 *Dec1^+/+^* and 11-16 *Dec1^-/-^* 2-3-month-old littermate mice; box plots indicate median and interquartile range. (**J**) GSIS from primary human pseudoislets transduced with Cas9-sgNT (control) or sg*DEC1* normalized to the first incubation (left). Insulin secretion fold change (16.7mM/2.8mM glucose) (middle). Total insulin content normalized to gDNA (right). Data are mean ±SEM from N =4 donors summarized in (L). (**K**) Change in relative expression of β-cell maturity markers in primary human pseudoislets transduced with sg*DEC1* relative to transduction with Cas9-sgNT. Dotted line indicates no change (1.0). Data are mean ±SEM from N =6 donors summarized in (L). (**L**) Summary of donor characteristics for primary human islet donors used in (**J-K**). (**M**) Left: heatmap shows the significance of correlations between mean β-cell *DEC1* expression level and insulin secretion under the indicated incubations in islets from non-diabetic (ND, n =25), autoantibody-positive (AAB+, n =10), type 1 diabetic (T1D, n =5), and type 2 diabetic (T2D, n =17) donors^75,119^. Right: scatter plot shows the correlation between mean β-cell *DEC1* expression and mean insulin secretion rate during 30 mM KCl stimulation in ND donors (Pearson’s r =0.44, p =0.028), with the blue line indicating the linear regression fit. *p <0.05, **p <0.01, ***p <0.001, t test.

**Extended Data Fig. 2.**
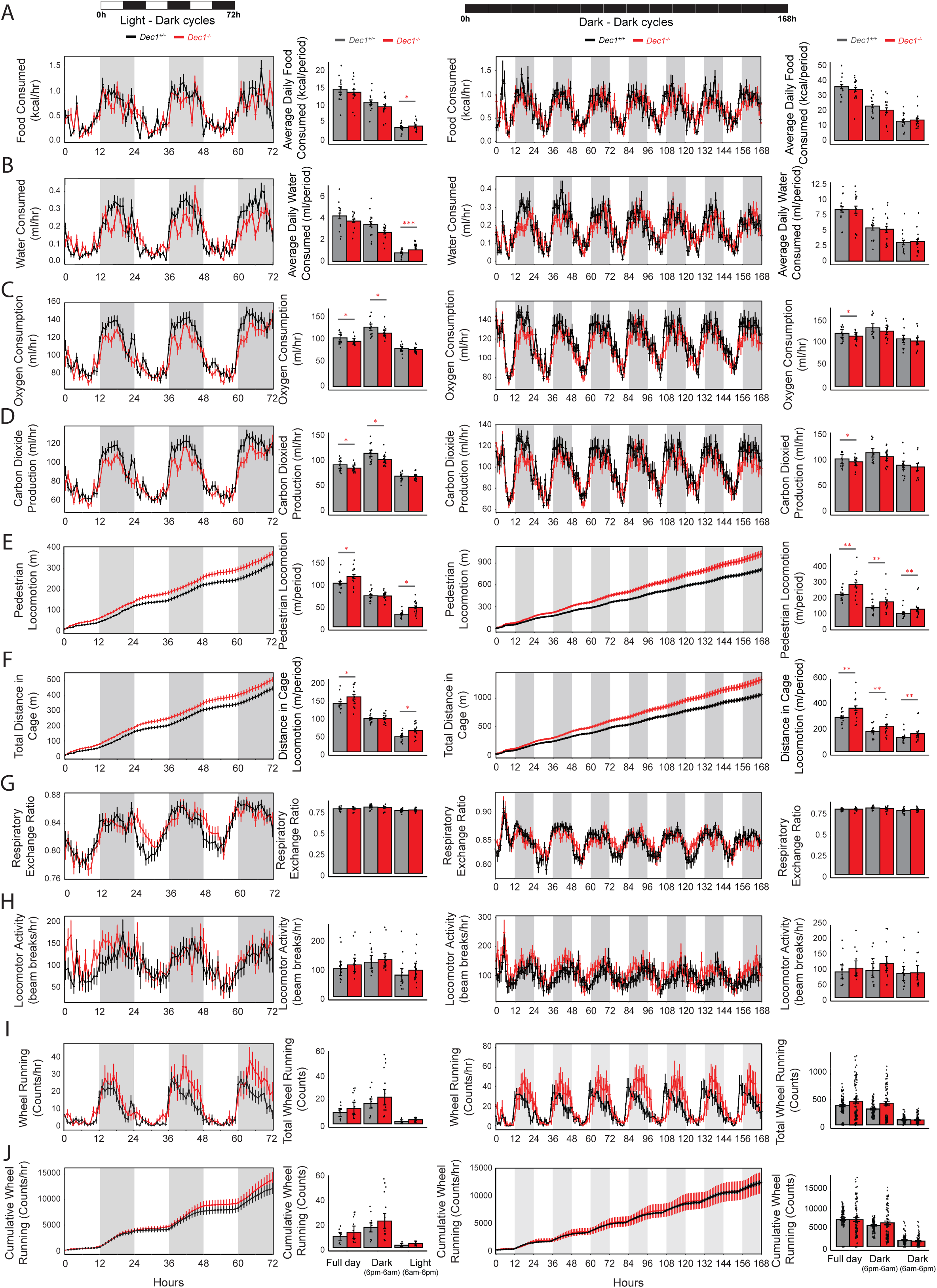
Metabolic and activity metrics of *Dec1*-knockout mice. **(A–J)** Metabolic and locomotor parameters measured in N = 13 *Dec1^+/+^*and N = 14 *Dec1^−/−^* 8-month-old littermate mice across 3 days in 12-hour light–dark (LD) cycles followed by 7 days in 12-hour dark–dark (DD) cycles. Parameters: food intake (A); water intake (B); oxygen consumption (C); carbon dioxide production (D); pedestrian locomotion (E); total distance traveled in cage (F); respiratory exchange ratio (G); locomotor activity (H); wheel running (I); cumulative wheel running (J). For each parameter, overall daily values were calculated by genotype and photoperiod. Data are presented as mean ± SEM. *p <0.05, **p <0.01, ANOVA test.

**Extended Data Fig. 3.**
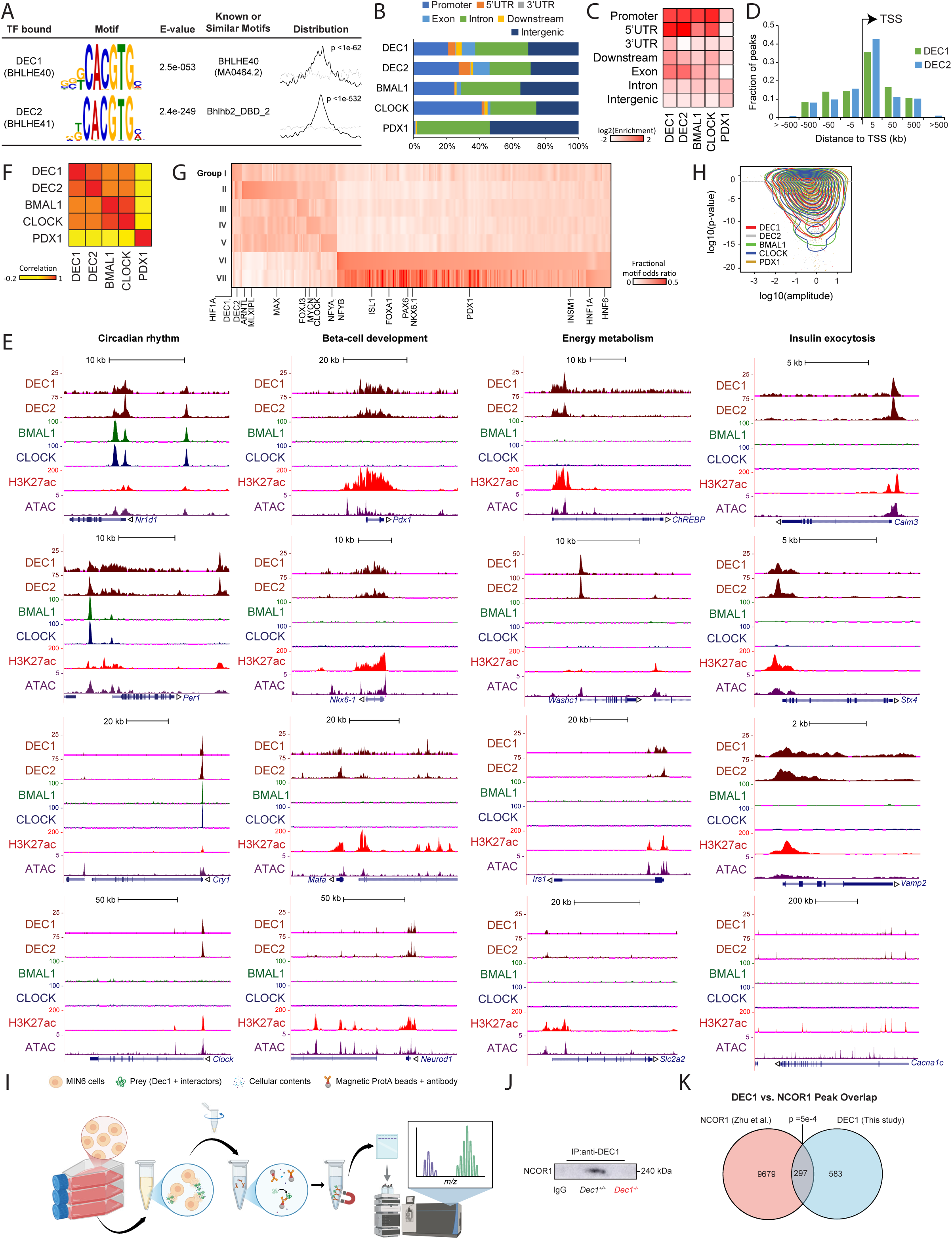
Circadian clock DEC regulators bind to energy metabolism and insulin secretion genes. (**A**) DEC1 and DEC2 binding peaks identified by ChIP-seq in β-cells enrich for the known DEC1 and DEC2 recognition motifs, respectively. Shown are the top motif identified by MEME-ChIP ^90^ across peak regions, its significance (E-value), top known motif match, and the distribution of the best matches to the motif across center-aligned peak regions. (**B**) Distribution of genomic locations for DEC1, DEC2, BMAL1, CLOCK, and PDX1 binding peaks from ChIP-seq in β-cells. (**C**) Relative enrichment for genomic locations among DEC1, DEC2, BMAL1, CLOCK, and PDX1 binding peaks from ChIP-seq in β-cells. (**D**) Distance to the nearest TSS for DEC1, DEC2, BMAL1, CLOCK, and PDX1 binding peaks from ChIP-seq in β-cells. (**E**) Genomic view of DEC1, DEC2, BMAL1, and CLOCK binding in β-cells at genes regulating circadian rhythm, beta-cell development, energy metabolism, and insulin exocytosis. Tracks display normalized ChIP and ATAC sequencing read density. Gene models are shown below. (**F**) Correlation between DEC1, DEC2, BMAL1, CLOCK, and PDX1 ChIP-seq signal, calculated across the union of their respective binding peaks. (**G**) Transcription factor motifs enriched across ChIP-seq binding clusters I to VII from Figure 3A. Heatmap shows relative enrichment across the groups for all motifs with group-specific expression. (**H**) Relationship between expression amplitude (peak-to-trough ratio), and significance of rhythmicity, as assessed by dryR^51^, for all genes linked to DEC1, DEC2, BMAL1, CLOCK, and PDX1 binding peaks from ChIP-seq in β-cells, indicating the p <0.05 threshold used. (**I**) Workflow for immunoprecipitation followed by mass spectrometry (IP-MS), created with BioRender.com. MIN6 (*Dec1^+/+^*or *Dec1^-/-^*) cells were cultured in flasks, collected and lysed to isolate their contents. Magnetic Protein A beads functionalized with anti-DEC1 (or anti-IgG control) antibody were incubated with cell lysate to pull down DEC1 and interacting/proximal proteins. Samples were then isolated by gel and loaded on a liquid-chromatography mass spectrometry instrument to ascertain relative abundance of protein species. (**J**) NCOR1 detected by immunoblot following DEC1 immunoprecipitation in MIN6 *Dec1^+/+^* but not *Dec1^-/-^* cells or following IgG immunoprecipitation in *Dec1^+/+^*MIN6 cells. (**K**) Intersection of DEC1 ChIP-seq peaks (This study) and NCOR1 ChIP-seq peaks in mouse liver (Zhu et al.)^85^. p-value, empirical test by regioneR^86^.

**Extended Data Fig. 4.**
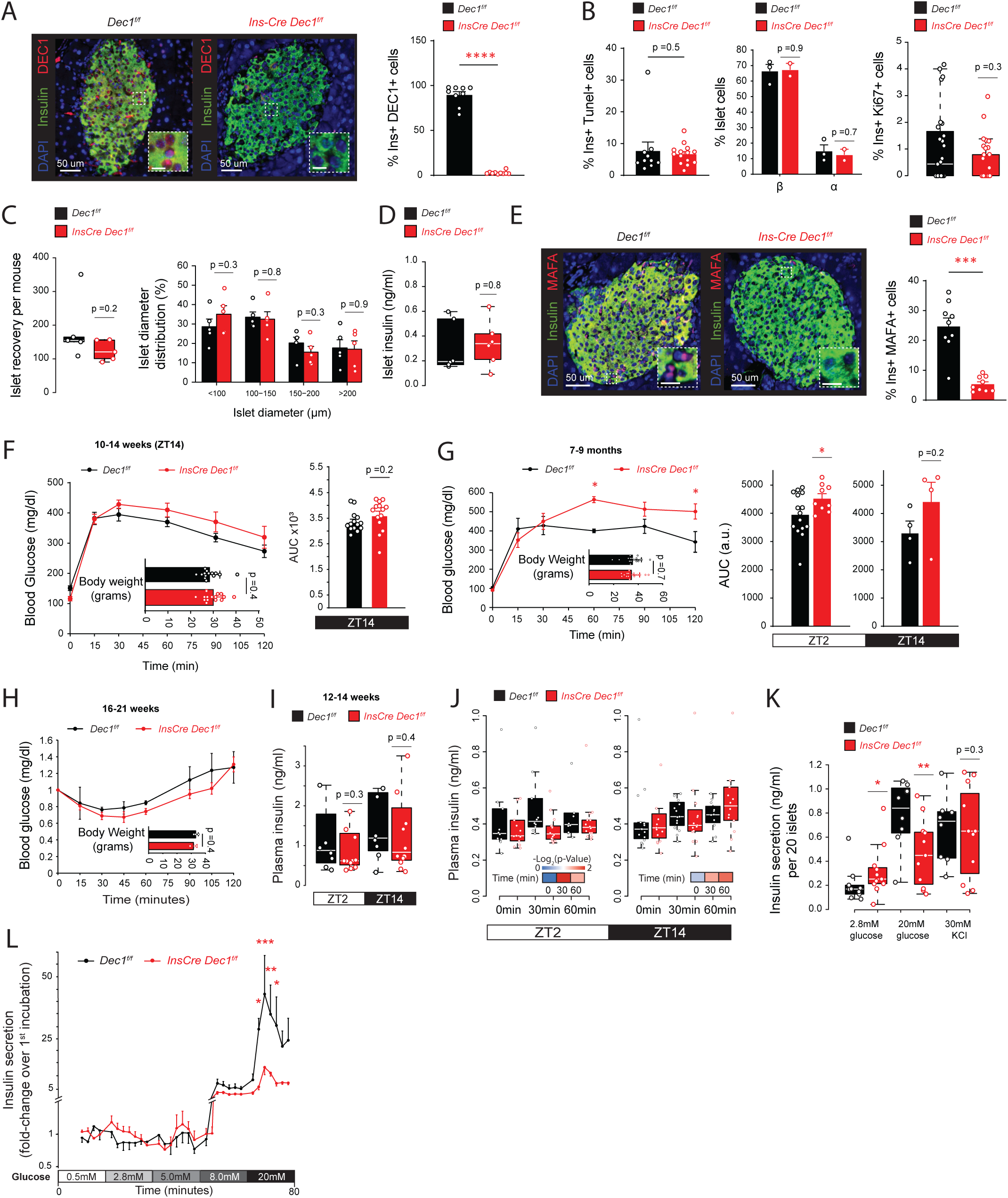
Glucose intolerance and immature islet function in young and aged β-cell specific *Dec1*-knockout mice despite normal weight, insulin sensitivity, and islet formation. (**A**) Left: immunofluorescence staining of insulin (green), glucagon (red), and DAPI (blue) in pancreata from N =2 *Dec1^f/f^* and 2 *InsCre Dec1^f/f^* 5-month-old littermate mice, acquired at 20x magnification. Insets scale bar =10 um. Right: fraction of DEC1-positive β cells. Data are mean ±SEM from n=9-10 islets. (**B**) Left: fraction of apoptotic (TUNEL+) β cells across pancreas sections from N =2 *Dec1^f/f^* and 2 *InsCre Dec1^f/f^* 5-month-old littermate mice; data are mean ±SEM. Center: fraction of α and β cells in islets isolated and pooled from N =2 *InsCre Dec1^f/f^* and 3 *Dec1^f/f^* 5-month-old littermate mice; data are mean ±SEM. Right: fraction of proliferating (Ki67+) β cells in islets isolated and pooled from N =2 *InsCre Dec1^f/f^*and 3 *Dec1^f/f^* 5-month-old littermate mice; boxplots indicate median and interquartile range. (**C**) Islets recovered per mouse (left), and their diameter distribution (right), from N =5 *InsCre Dec1^f/f^*and 5 *Dec1^f/f^* 7-9-month-old littermate mice. (**D**) Insulin content of islets isolated and pooled from N =6 *InsCre Dec1^f/f^* and 6 *Dec1^f/f^*5-month-old littermate mice. Data are from n =6 independent measurements of 20 size-matched islets. (**E**) Left: immunofluorescence staining of insulin (green), MAFA (red), and DAPI (blue) in N =2 *Dec1^f/f^* and 3 *InsCre Dec1^f/f^* littermate 5-month-old littermate mice, acquired at 20x magnification. Insets scale bar =10 um. Right: fraction of MAFA-positive β cells. Data are mean ±SEM of isolated islets pooled from N =2 *InsCre Dec1^f/f^* and 3 *Dec1^f/f^*5-month-old littermate mice from n=5-7 replicate measurements. (**F**) Glucose tolerance in 14-hour fasted N =14 *InsCre Dec1^f/f^* and 15 *Dec1^f/f^* 10-14-week-old littermate mice following intraperitoneal glucose administration at ZT14. Data are mean ±SEM. (**G**) Glucose tolerance in 14-hour fasted N =16 *InsCre Dec1^f/f^* and 17 *Dec1^f/f^*7-9-month-old littermate mice following intraperitoneal glucose administration at ZT2. Data are mean ±SEM. (**H**) Insulin tolerance in 4-hour fasted N =3 *InsCre Dec1^f/f^* and 3 *Dec1^f/f^*16-21-week-old littermate mice following intraperitoneal insulin administration at ZT2. Data are mean ±SEM. (**I**) Plasma insulin in *ad libitum* fed N =12 *InsCre Dec1^f/f^* and 8 *Dec1^f/f^*3-month-old littermate mice at ZT2 and ZT14. Data are mean of n =2 replicate measurements. (**J**) Plasma insulin in mice from Fig. 4C at 0, 30, and 60 minutes following intraperitoneal glucose administration at ZT2 and at ZT14. (**K**) Glucose-stimulated insulin secretion of isolated islets from N =6 *InsCre Dec1^f/f^* and 6 *Dec1^f/f^* 8-month-old littermate mice, each with n =2 replicate measurements of 20 size-matched islets under the indicated sequential static incubations. (**L**) Insulin secretion dynamics in perifused islets from N =2 *InsCre Dec1^f/f^* and 2 *Dec1^f/f^*8-month-old littermate mice, each with n =3 replicate measurements of 20 size-matched islets under increasing glucose concentrations. Data are mean ±SEM, normalized to the mean of the first incubation. *p <0.05, **p <0.01, ***p <0.001, ****p <0.0001, t test.

**Extended Data Fig. 5.**
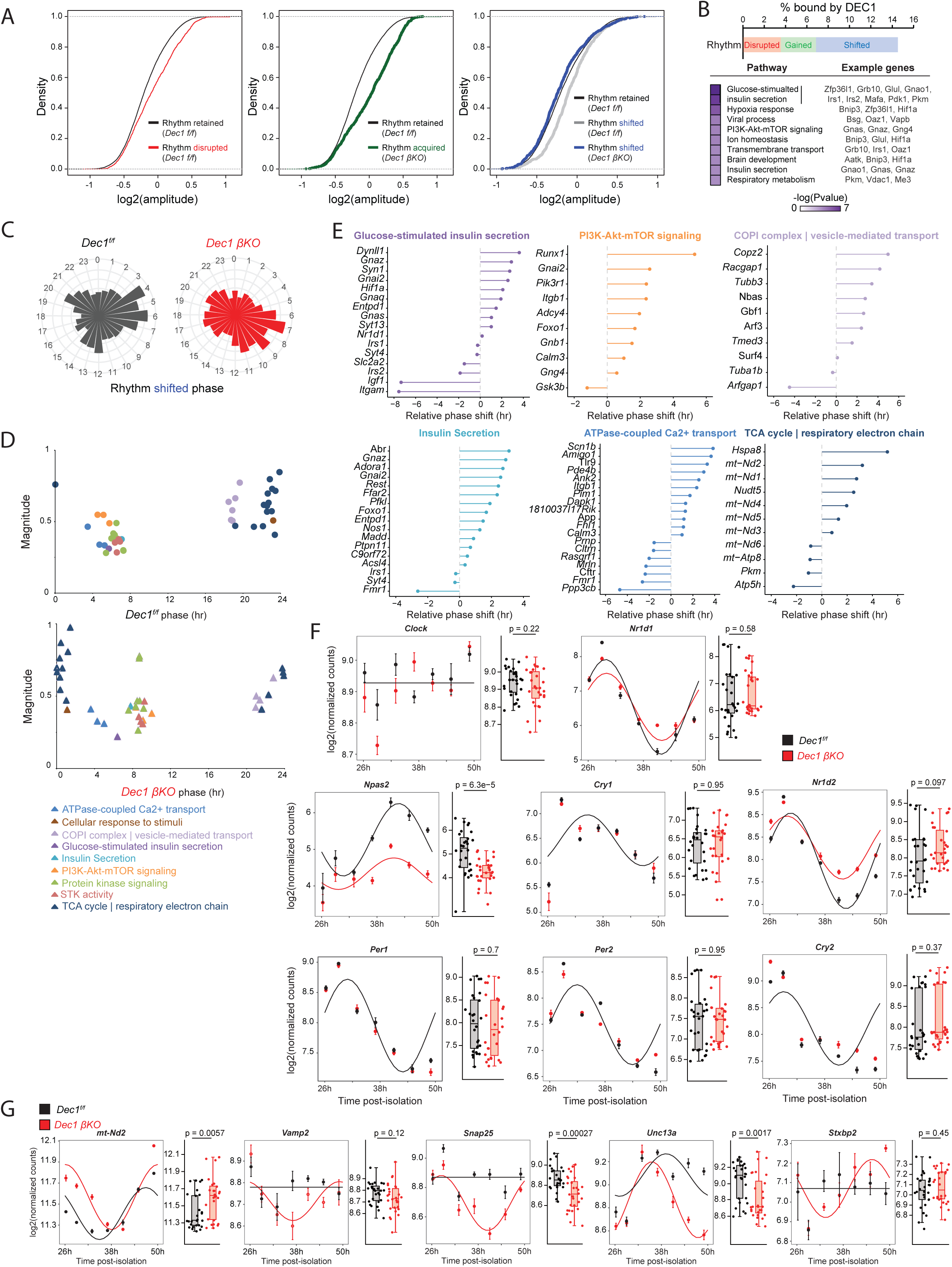
DEC1 coordinates β-cell respiratory metabolism and insulin secretion gene rhythms. (**A**) Distribution of rhythm amplitudes for genes whose expression rhythm is disrupted (left), enhanced (middle), and shifted (right) in *Dec1 βKO* islets, compared to those of rhythm-retaining genes. (**B**) Among genes with rhythmic expression that is disrupted, enhanced, or shifted in *Dec1 βKO* islets, the fraction that is bound by DEC1 in β cells is shown at the top. Pathways enriched among the genes disrupted, enhanced, or shifted in *Dec1 βKO* islets that are bound by DEC1 are shown below. (**C**) Phase distribution of transcripts whose expression rhythm is shifted in *Dec1 βKO* islets. (**D**) Average magnitude and phase of expression of gene sets driving enrichment for the indicated biological themes among genes whose expression rhythm is shifted in *Dec1 βKO* islets, colored by biological theme. (**E**) Relative phase shift, defined as the shortest angular difference, for genes whose expression rhythm is shifted in *Dec1 βKO* islets, grouped and colored by biological theme. (**F-G**) Examples of genes with unaltered (F) and altered (G) temporal expression patterns in *Dec1 βKO* islets. Data are mean from n =4 replicate measurements of islets isolated and pooled from N =2 *Dec1*-*βKO* and 2 *Dec1^f/f^*5-month-old littermate mice. p-value, t test.

**Extended Data Fig. 6.**
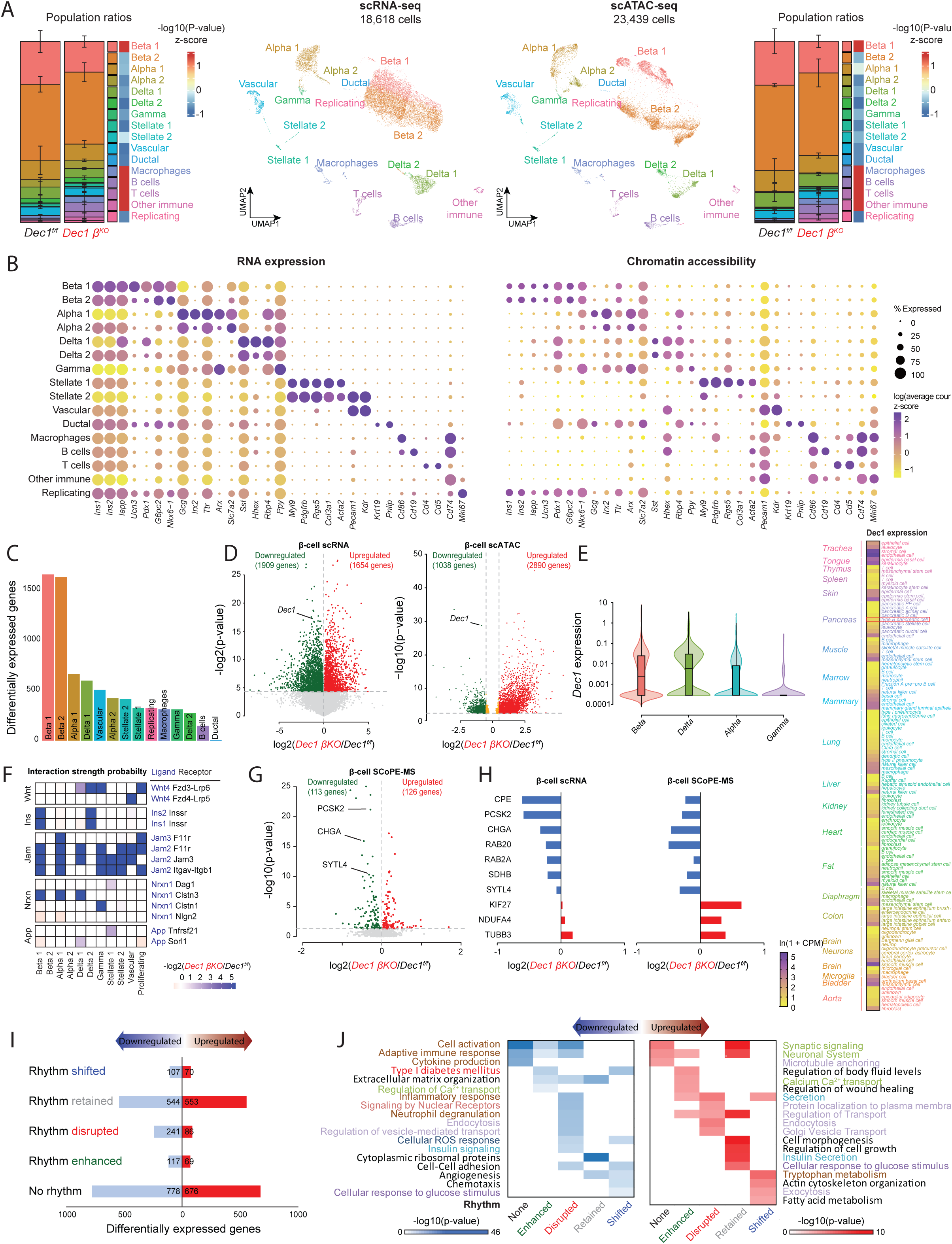
Altered chromatin, transcription, and protein level regulation of glucose metabolism and glucose-dependent insulin secretion in β-cells of *Dec1-βKO* mice. (**A)** Single-cell clustering by gene expression profiles (left, scRNA-seq) and by open chromatin profiles (right, scATAC-seq) in islets isolated from N =2 *Dec1*-*βKO* and 2 *Dec1^f/f^* 7-month-old littermate mice, each sequenced separately, with population ratios and the relative significance of ratio changes (hypergeometric test) shown to the sides. (**B**) Quantification of RNA (left) and open chromatin (right) reads mapping to selected marker genes for each cell cluster in (A). Color represents z-normalized average read counts, and diameter denotes fractional expression. (**C**) Distribution of genes that are differentially expressed in the indicated cell types in islets isolated from either or both of N =2 *Dec1-βKO* relative to 2 *Dec1^f/f^* 7-month-old littermate mice, each sequenced separately. (**D**) Changes in gene expression (left) and chromatin accessibility (right) in β-cells pooled from N =2 *Dec1-βKO* relative to 2 *Dec1^f/f^* 7-month-old littermate mice, each sequenced separately. **(E)** *Dec1* expression across islet cell types and mouse tissues. Left: violin plots show *Dec1* expression across β, δ, α, and γ cells in islets from (A). Right: Heatmap shows *Dec1* expression across mouse tissues and cell types from the Tabula Muris project^74^, derived from N = 4 wild-type mice, highlighting pancreatic β cells (red box).(**F**) Changes in intercellular ligand-receptor interaction strength probabilities (see Methods) in islets pooled from N =2 *Dec1-βKO* relative to 2 *Dec1^f/f^* 7-month-old littermate mice, for 7 of the top 10 differentially active interactions where the ligand expression is significantly altered in Beta 1 cells from (A) with higher expression of insulin and maturity markers, based on scRNA-seq data. Grouped ligand-receptor pairs are indicated to the right. (**G-H**) Proteome changes in β-cells pooled from N =2 *Dec1-βKO* relative to 2 *Dec1^f/f^* 8-month-old littermate mice, each run separately (G). scRNA and protein changes in β-cells for selected targets are shown in (H). (**I**) Distribution of genes differentially expressed in β-cells pooled from N =2 *Dec1*-*βKO* relative to 2 *Dec1^f/f^* 7-month-old littermate mice, each run separately, among non-rhythmic or enhanced, disrupted, retained, or shifted rhythmicity dryR models. (**J**) Pathways enriched among non-rhythmic genes or genes with enhanced, disrupted, retained, or shifted rhythmicity that are downregulated (left) or upregulated (right) in β-cells pooled from N =2 *Dec1*-*βKO* relative to 2 *Dec1^f/f^* 7-month-old littermate mice, each run separately.

**Extended Data Fig. 7.**
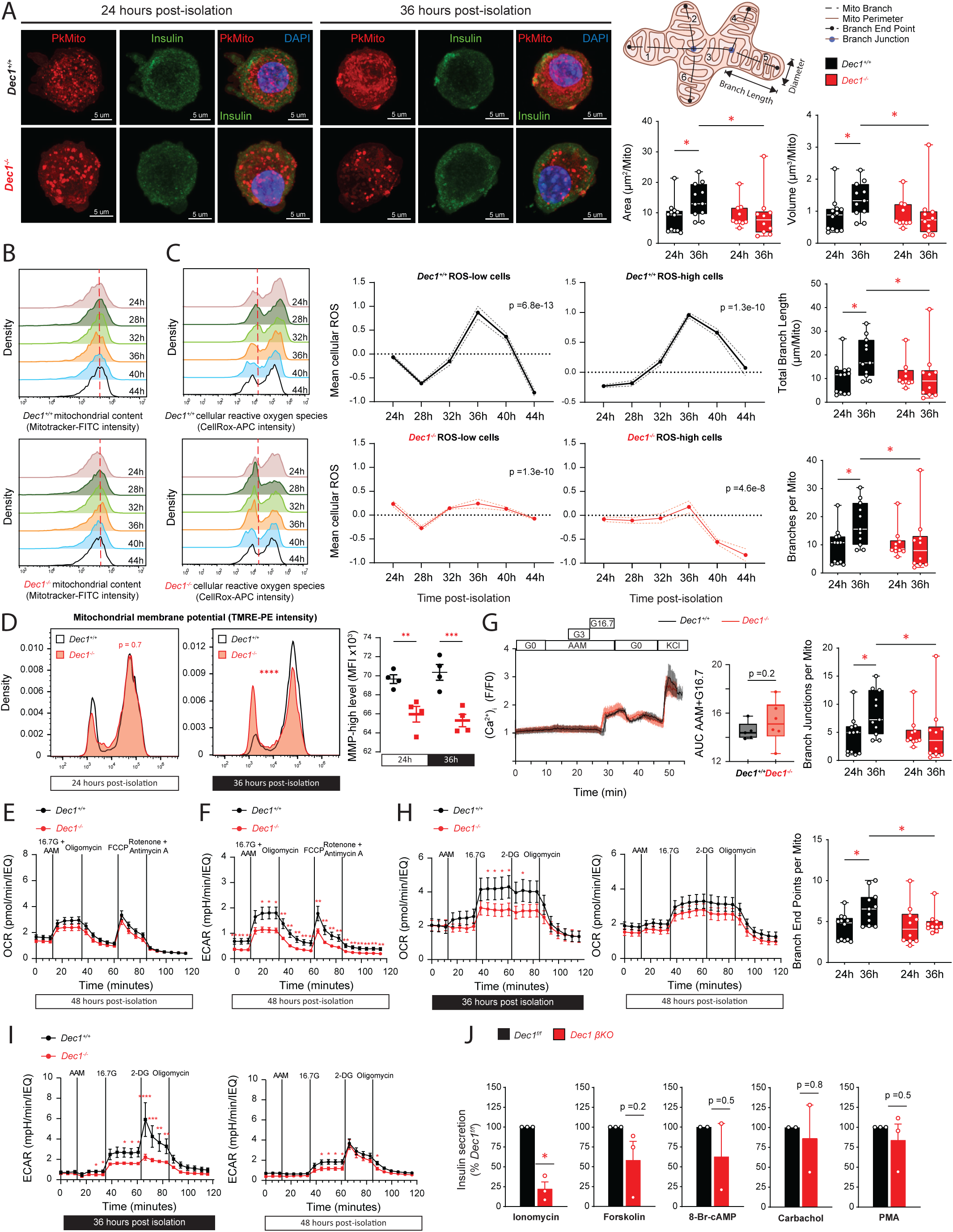
Disruption of circadian mitochondrial dynamics and energy metabolism underlies immature β-cell function upon DEC1 loss. (**A**) Left: immunofluorescence staining of mitochondria (red; PKmito Orange FX), insulin (green), and DAPI (blue) in β-cells isolated from *Dec^+/+^* and *Dec1^-/-^*mice at 24h and 36h post-isolation, acquired at 63x magnification. Center: diagram skeletonizes a mitochondrion to mark 6 branches (1-6), 5 branch end points, and 2 branch junctions, and shows the perimeter used to calculate volume and sphericity. Quantifications of the indicated structural features at 24h and 36h are shown to the right and below. Data are distributions of confocal imaging measurements of β-cells isolated and pooled from N =3 *Dec^+/+^* and 2 *Dec1^-/-^* 2-3-month-old littermate mice with n =10-12 independent fields of view, each with >50 mitochondria. (**B**) Mitochondrial content across islet-cells isolated from N =2 *Dec1^+/+^* and 2 *Dec1^-/-^* 2-4-month-old littermate mice, assayed by MitoTracker Green fluorescence intensity at 4h intervals for 20h. (**C**) Cellular reactive oxygen species (ROS) distributions (left) and their 24h rhythmicity analysis (middle and right) in islet-cells isolated from N =2 *Dec1^+/+^* and 2 *Dec1^-/-^*2-4-month-old littermate mice. Distributions display flow cytometry of CellRox Deep Red intensity in islet-cells-cells analyzed every 4h for 20h. The separation between ROS-low and-high populations is indicated by the red dotted line in the left diagrams. Rhythmicity analysis quantifies detrended (baseline-subtracted) mean ± SEM intensity for ROS-low and-high cell populations separately. Rhythmicity p-value, RAIN. (D) Mitochondrial membrane potential (MMP) across islet-cells isolated from N =2 *Dec1^+/+^* and 2 *Dec1^-/-^* 2-4-month-old littermate mice (left and center), determined by TMRE staining at 24h and 36h post-isolation, and quantification of mean fluorescence intensity (MFI) for MMP-high cells (right). Data are distributions and mean values from n=4 replicate measurements, each with >5000 live singlet cells. (**E-F**) Metabolic flux in islets isolated from N =5 *Dec1^+/+^* and 5 *Dec1^-/-^* 2-4-month-old littermate mice quantified from measurements of oxygen consumption rate (OCR) (E) and extracellular acidification rate (ECAR) (F) in response to sequential incubations with 16.7mM glucose and 3.5mM amino acid mixture (16.7G + AAM), oligomycin, FCCP, and to rotenone and antimycin A at 48h post-isolation. Data are mean ±SEM from n =4-6 replicate measurements of 12 similar-sized islets. (**G**) Ca^2+^ influx measured in islets isolated from N = 2 *Dec1^+/+^* and 2 *Dec1^-/-^*2-3-month-old mice with n =5-7 independent fields of view quantified by Fura-2 staining in response to sequential incubations with no glucose (G0), a 3.5mM amino acid mixture (AAM), AAM supplemented with 3mM glucose (G3), AAM supplemented with 16.7mM glucose (G16.7), no glucose, and 30mM KCl, with influx during AAM+G16.7 quantified to the right. (**H-I**) Metabolic flux in islets isolated from N =7 *Dec1^+/+^* and 5 *Dec1^-/-^* 2-4-month-old littermate mice quantified from measurements of oxygen consumption rate (OCR) (H) and extracellular acidification rate (ECAR) (I) in response to sequential incubations with 3.5mM amino acid mix, 16.7mM glucose, 50mM 2-DG, and oligomycin at 36h and 48h post-isolation. Data are mean ±SEM from n =4-7 replicate measurements of 12 similar-sized islets. (**J**) Glucose and secretagogue-stimulated insulin secretion fold-change (20mM / 2.8mM glucose) in sequential static incubations of size-matched islets isolated from N =4-5 *Dec1*-βKO and 4-6 *Dec1^f/f^* 7-month-old littermate mice, each with n =2-3 replicate measurements, normalized to intracellular insulin content. Data are mean ±SEM. *p <0.05, **p <0.01, ***p <0.001, ****p <0.0001, t-test [(A), (D) (right), (G), (J)], chi-squared test (D) (left), two-way ANOVA [(F), (H), (I)].

**Extended Data Fig. 8.**
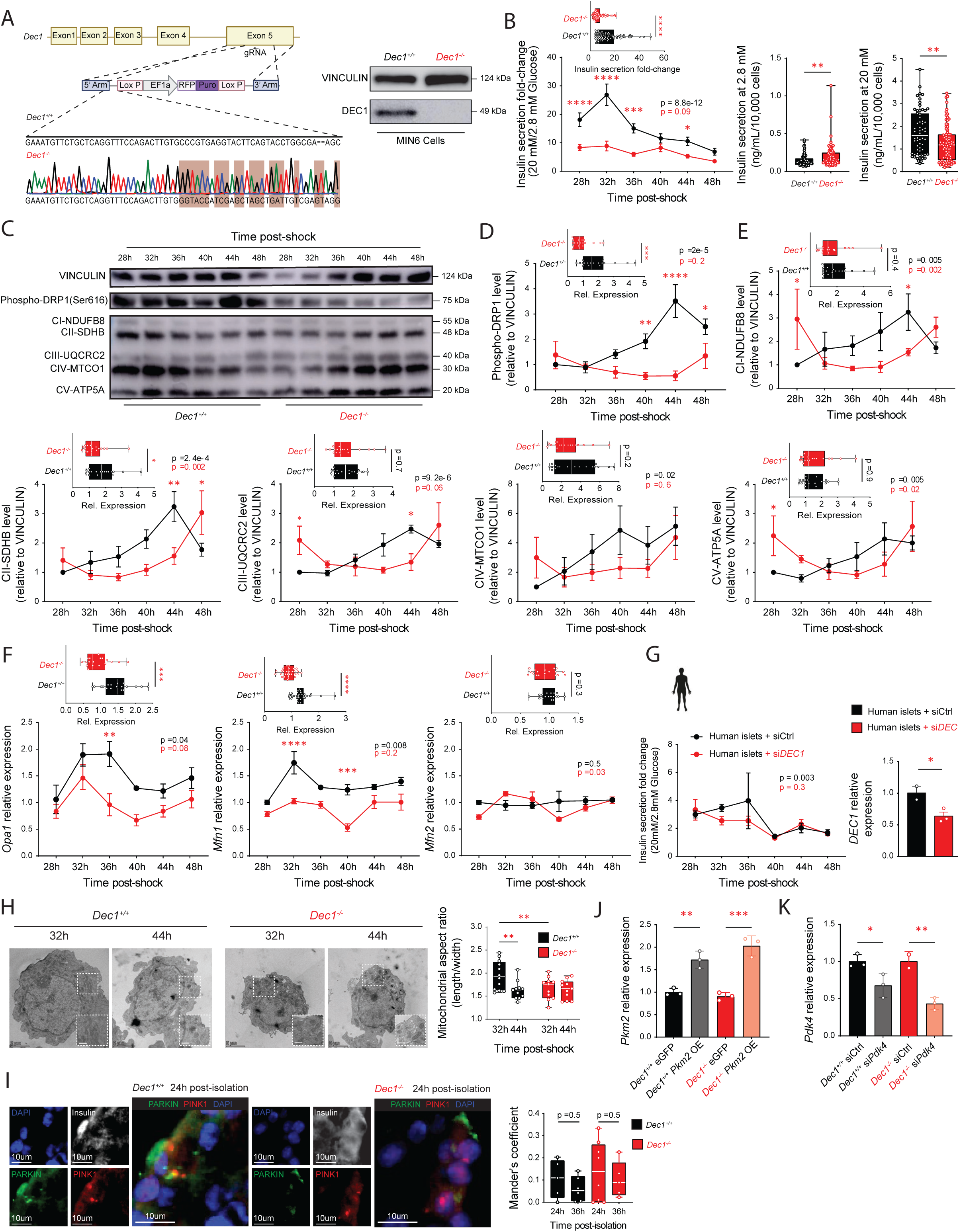
Molecular basis for disrupted circadian mitochondrial remodeling and metabolic immaturity in *Dec1*-knockout β-cells. (**A**) Validation of *Dec1* knockout in MIN6 cells. Guide RNAs targeting *Dec1*’s exon 5 and a selectable donor template were introduced into MIN6 cells (see Methods). Sanger sequencing traces across the targeted region confirms disruption of the endogenous sequence. Western blotting verifies loss of DEC1 protein in edited cells relative to mock-treated control cells. (**B**) Insulin secretion fold-change (20 mM / 2.8 mM glucose) sampled every 4h for 20h in media shock-synchronized *Dec1^+/+^* and *Dec1^−/−^* MIN6 cells (left). Insulin secretion at 2.8mM glucose (center) and 20mM glucose (right). Data are mean ± SEM. Rhythmicity p-value, RAIN. (**C**) Immunoblot analysis of mitochondrial fission and respiratory chain proteins every 4h for 20h in media shock-synchronized *Dec1^+/+^* and *Dec1^−/−^* MIN6 cells. Proteins quantified were phospho-Ser616 DRP1, NDUFB8 (Complex I), SDHB (Complex II), UQCRC2 (Complex III), MTCO1 (Complex IV), and ATP5A (Complex V), with VINCULIN as loading control. (**D-E**) Quantification of phospho-DRP1 (D) and OXPHOS complex I to V proteins (E) across *Dec1*^+/+^ and *Dec1^−/−^* MIN6 cells from (C). Data are mean ± SEM. Rhythmicity p-value, RAIN. (**F**) Expression of mitochondrial fusion genes (*Opa1*, *Mfn1*, and *Mfn2*) every 4h for 20h in media shock-synchronized *Dec1^+/+^* and *Dec1^−/−^*MIN6 cells. Data are mean ± SEM. Rhythmicity p-value, RAIN. (**G**) Insulin secretion fold change (20 mM / 2.8 mM glucose) every 4h for 20h in forskolin shock-synchronized primary human islets transfected with control siRNA (siCtrl) or siRNA targeting *DEC1* (si*DEC1*) (left). Relative *DEC1* expression confirming knockdown (right). Data are mean ± SEM. Rhythmicity p-value, RAIN. (**H**) Transmission electron microscopy images of media shock-synchronized *Dec1^+/+^* and *Dec1^−/−^* MIN6 cells (left) at 32h and 44h post-shock, acquired at 5300x (overview) and 17500x (insets) magnification. Quantification of mitochondrial aspect ratio (length/width) (right). Data are mean ± SEM with n =11-12 replicate measurements. Insets scale bar is 250 nm. (**I**) Immunofluorescence stainning of insulin (green), PARKIN (red), PINK1 (white), and DAPI (blue) in islets isolated and pooled from N =2 *Dec1^+/+^* and 3 *Dec1^−/−^*2-month-old littermate mice (left), acquired at 20x magnification. Quantification of PARKIN and PINK1 co-localization by Mander’s coefficient at 24h and 36h post-isolation (right). Data are distribution of n =5-8 replicate measurements. (**J**) *Pkm2* relative expression in islets isolated and pooled from N =6 *Dec1^+/+^* and 5 *Dec1^−/−^* 2-3-month-old littermate mice and transduced with *Pkm2*-overexpressing (OE) or eGFP control plasmids. Data are mean ± SEM with n =3 replicate measurements. (**K**) *Pdk4* relative expression in islets isolated and pooled from N =3 *Dec1^+/+^* and 3 *Dec1^−/−^* 2-3-month-old littermate mice and transfected with *Pdk4*-targeing siRNA (si*Pdk4*) or control siRNA (siCtrl). Data are mean ± SEM with n =2-3 replicate measurements. *p <0.05, **p <0.01, ***p <0.001, ****p <0.0001, t-test {insets[(C), (E), (F), (G), (I)], (H)}, two-way ANOVA [(C), (E), (F), (G)], one-way ANOVA [(J), (K), (L)]

